# Extracellular Polymeric Substances Drive Curvibacter Host Adaptation and Co-Speciation with Hydra

**DOI:** 10.1101/2025.09.08.674843

**Authors:** Lukas Becker, Liam Fürbach, Timo Minten, Jay Bathia, Marius Karbach, Sylvain Foret, Nicolas M. Winterfeldt, Markus Pauly, Ilka M. Axmann, Sebastian Fraune

## Abstract

The interactions between hosts and their microbial symbionts play a crucial role in shaping biological diversity and ecosystem function. Bacteria can adapt to specific host environments over evolutionary timescales, leading to co-speciation and the formation of specialized host-microbe relationships. Understanding the molecular mechanisms underlying these adaptations provides key insights into the evolution of symbiosis and the stability of microbial communities. This study investigates the co-speciation and molecular adaptations of *Curvibacter* to its host *Hydra*, a well-established model for the study of host-microbe interactions. We provide strong evidence of co-speciation, as demonstrated by phylogenetic congruence between different *Hydra* species and their corresponding *Curvibacter* symbionts, along with preferential recolonization of germ-free *Hydra* by their native *Curvibacter* strains. Comparative genomic analyses reveal that host-associated *Curvibacter* strains exhibit distinct metabolic and biosynthetic adaptations compared to their free-living relatives. Specifically, the enrichment of proteins involved in sugar metabolism and transport, as well as the selective purification of proteins linked to macromolecule biosynthesis, highlights the specialization of *Curvibacter* symbionts within the *Hydra* glycocalyx. Functional experiments identify a symbiont-specific extracellular polymeric substances (EPS) operon as key factor for microbial adhesion and host colonization, underscoring its role in facilitating symbiont specificity and stability. These findings provide insights into the molecular mechanisms driving host-microbe co-evolution and highlight the evolutionary forces shaping microbial specialization within host-symbiont relationships.

**Significant Statement:** The vast majority of animal and plant species are associated with microbial organisms that offer a wide range of benefits for development, physiology, and health. Understanding the evolutionary forces that shape these symbiotic interactions is crucial for elucidating the mechanisms through which organisms mitigate environmental stress and enhance their resilience, particularly in response to rapidly changing environmental conditions. Using advanced bioinformatic, bioanalytical, and genetic approaches, this study demonstrates that *Curvibacter* symbionts have evolved specific extracellular polymeric substances to enhance their ability to form a specific symbiosis with *Hydra*.

## Introduction

Microbial communities that colonize host organisms enhance the fitness of the host by supporting a range of functions, including the provision of nutrients, the maturation of the immune system and the development of resistance to pathogens (Fraune & Bosch, 2010; Mcfall-Ngai et al., 2013). In return, microbial communities benefit from the ecological niche provided by the host (Kopac & Klassen, 2016; Obeng et al., 2021). Consequently, the host and its associated microbiota constitute a complex, integrated biological entity, designated as ‘holobiont’ or ‘metaorganism’ (Bosch & McFall-Ngai, 2011; Rosenberg & Zilber-Rosenberg, 2018; Zilber-Rosenberg & Rosenberg, 2008), which is shaped by evolutionary pressures that drive mutual adaptation and specialization over time. Studying the underlying evolutionary processes is critical for understanding of how bacteria affect metaorganism maintenance and fitness. The phenomenon of phylosymbiosis, whereby the similarity of associated microbial communities correlates with the evolutionary history of their hosts, has been documented across diverse host clades (Brooks et al., 2016; Franzenburg et al., 2013; Hayward et al., 2021; Lim & Bordenstein, 2020). This observation implies that host evolution exerts a long-term influence on the composition of their symbiont communities. However, the extent to which shared environments or co-evolutionary processes between host and symbiont determine the observed microbial patterns remains unclear in most cases (Moran & Sloan, 2015).

Due to its simplicity, we use the freshwater polyp *Hydra* and its symbiont *Curvibacter* as host-microbe model to study mechanisms of symbiont adaptation and specificity (Minten-Lange & Fraune, 2020). *Hydra* is a small freshwater polyp belonging to the phylum *Cnidaria*, a sister group to all bilaterians. *Hydra* features a relatively simple body plan and lifestyle, facilitating efficient experimental analysis. Comparing the bacterial communities of different *Hydra* species maintained in the lab revealed a high degree of pyhlosymbiosis (Franzenburg et al., 2013). In addition, *Hydra* polyps sampled from the field are associated with similar bacterial communities as polyps from the laboratory (Fraune & Bosch, 2007; Taubenheim et al., 2022). These associated microbial species are located either epibiotic within the glycocalyx of *Hydra* or endosymbiotic within the epithelial cells (Fraune et al., 2015; Fraune & Bosch, 2007). The innate immune system of *Hydra* relies on a rich repertoire of antimicrobial peptides and an evolutionary conserved set of pattern recognition receptors (Augustin et al., 2009, 2017; Bosch et al., 2009; Franzenburg et al., 2012; Franzenburg et al., 2013; Fraune et al., 2010; Jung et al., 2009; Klimovich & Bosch, 2024). This innate immune repertoire protects the metaorganism *Hydra* against pathogens and maintains a homeostasis with beneficial microbes. Conversely, the *Hydra* microbiota has multiple effects on the host. Polyps with their microbiota experimentally removed (germ-free animals) exhibit behavioral changes (Murillo-Rincon et al., 2017), and are prone to fungal infection (Fraune et al., 2015). In addition, members of the microbiome can induce tumor development (Boutry et al., 2023; Rathje et al., 2020), asexual reproduction (Rahat & Dimentman, 1982) and pattern formation which is mediated via the activation of host peptides antagonizing the Wnt signaling pathway (Taubenheim et al., 2020). The most abundant bacterial colonizer of *Hydra vulgaris* is *Curvibacter*, a Betaproteobacterium belonging to the family of *Comamonadaceae* (Fraune et al., 2015). *Curvibacter* colonizes the mucus-like layer of the ectodermal epithelial cells of *Hydra* together with several other microbial colonizers (Fraune et al., 2015). *Curvibacter* is involved in antifungal activity (Fraune et al., 2015) and induces the production of the Eco host peptides, which act as Wnt antagonists and thereby influence the pattern formation and behavior of stem cells (Taubenheim et al., 2020). To sustain long-term bacterial functions vertical transmission of beneficial bacterial cells is crucial. In *Hydra*, *Curvibacter* is transferred during asexual reproduction by budding, whereby the bacterial cells passively migrate from the mother polyp to the bud (Franzenburg et al., 2013; Fraune et al., 2010). During embryogenesis, however, *Curvibacter* is absent in early stages, due to the antimicrobial peptide Periculin but colonizes the embryo in later stages, likely from the mother’s tissue to the embryo’s cuticle (Fraune et al., 2010). After hatching, *Curvibacter* attaches from the eggshell to newly hatched polyps, ensuring transmission to the next sexual generation (Franzenburg et al., 2013; Minten-Lange & Fraune, 2020). In addition, *Curvibacter* can effectively colonize germ-free *Hydra* polyps when added to the surrounding medium (Fraune et al., 2015; Wein et al., 2018). This demonstrates that *Curvibacter* has maintained the ability to have a biphasic life cycle in which it can alternate between a host-associated and a free-living phase.

In this study, we address the specificity of different *Curvibacter* strains for their respective *Hydra* hosts. We observed strong congruent phylogenetic patterns between *Curvibacter* and its *Hydra* hosts, supporting the presence of co-speciation (Kawaida et al., 2013). Reciprocal recolonization experiments with three different *Hydra* species and their associated *Curvibacter* strains showed that all strains performed best in recolonizing their native *Hydra* hosts, emphasizing the evidence for co-speciation. To further investigate host adaptation within the *Curvibacter* genus, a comparative genomic analysis of free-living and host-associated species was performed, which revealed greater genetic similarity among host-associated *Curvibacter* strains compared to their free-living relatives. This analysis also identified an extracellular polymeric substances (EPS) operon more prevalent in host-associated *Curvibacter* strains. Knockout mutants lacking components of the EPS operon (Δ*epsH* and Δ*wcaJ*) exhibited altered levels of three EPS-associated monosaccharides and reduced ability to recolonize host tissue. The findings indicate that *Curvibacter* has evolved specific adaptations to thrive within the *Hydra* host species, facilitated by exopolysaccharides that reinforce the symbiotic relationship with the host.

## Results

### Co-speciation of *Curvibacter* and *Hydra*

Recognizing that *Curvibacter* is a consistent part of the microbiome of many *Hydra* species (Franzenburg et al., 2013), we performed a cultivation effort to isolate *Curvibacter* strains from all species available in the laboratory. In total, this isolation procedure yielded 16 additional *Curvibacter* strains from six different *Hydra* species (**Figure 1A**). To investigate potential co-speciation between *Curvibacter* and *Hydra*, we compared the phylogeny of the six *Hydra* species (**Figure 1A, left**) with those of 16 corresponding *Curvibacter* strains and the already described strain Curvibacter AEP 1.3 (Pietschke et al., 2017) as well as the *Curvibacter* strain co-sequenced with the *Hydra magnipapillata* genome (NCBI:txid667019) (**Figure 1A, right**). Phylogenetic analysis of the 16S rRNA nucleotide sequences from *Curvibacter* revealed six distinct clusters, each of which aligns congruently with its respective *Hydra* host, as reflected in the phylogeny of the cytochrome C oxidase subunit I gene of the six *Hydra* species. The closely related *Curvibacter* species within each of the six clusters originate from the same *Hydra* species. This high congruency suggests a co-speciation of *Curvibacter* and *Hydra* since early *Hydra* evolution.

**Figure 1:**
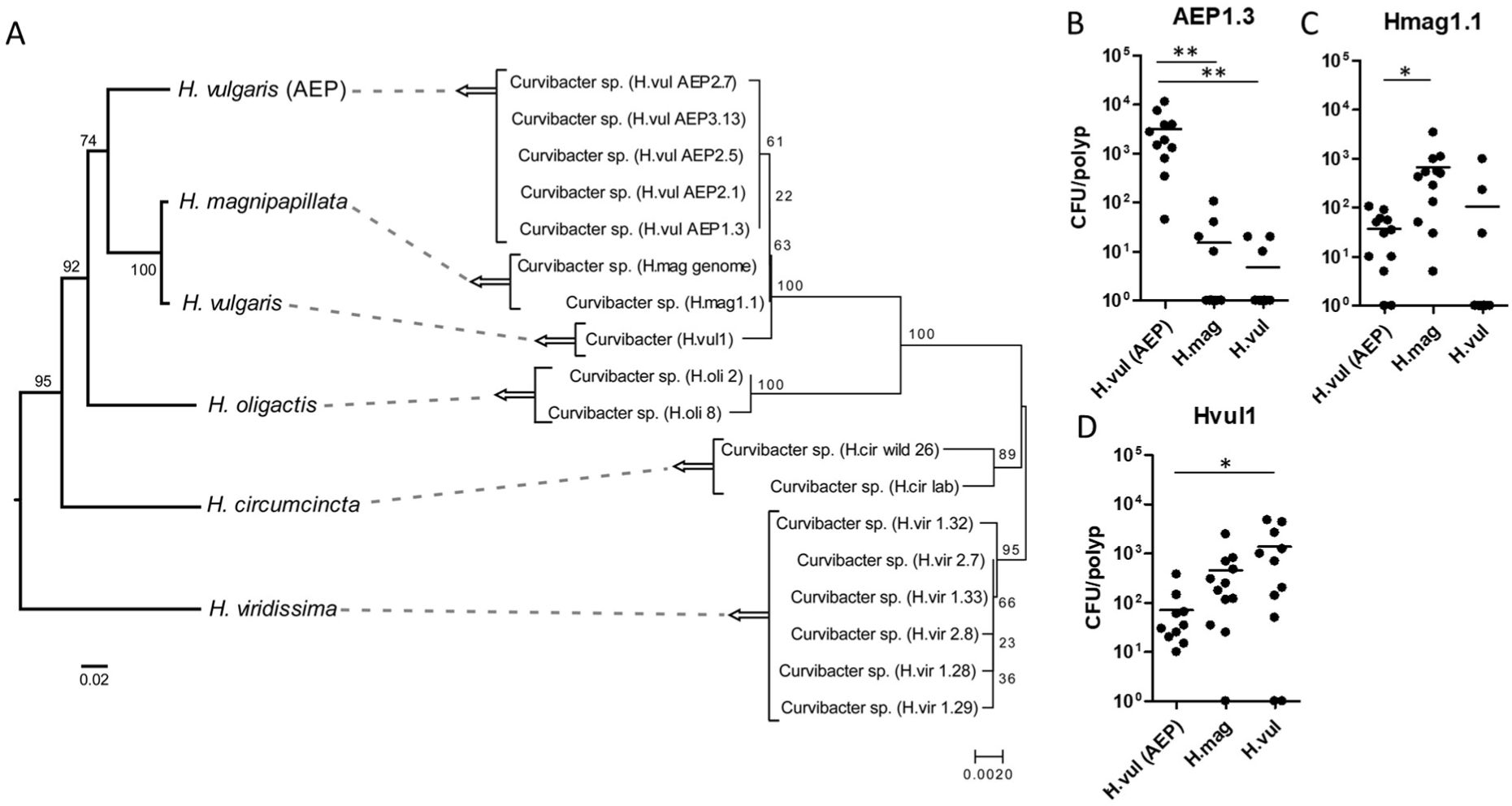
Co-speciation of *Hydra* and their associated *Curvibacter* strains. (**A**) Comparison of the phylogenetic trees of *Hydra* (left) and the associated *Curvibacter* species (right). (*Left*) Phylogenetic tree of *Hydra* species based on cytochrome oxidase genes (maximum likelihood, general time reversible (GTR+I)). Bootstrap values are shown at the corresponding nodes. The branch-length indicator displays 0.02 substitutions per site. *(Right)* Phylogenetic tree of *Curvibacter* isolates based on the 16S rRNA genes (maximum likelihood, Hasegawa-Kishino-Yano model). Bootstrap values are shown at the corresponding nodes. The branch-length indicator displays 0.002 substitutions per site. Note: the phylogenies of the *Hydra* hosts and their associated *Curvibacter* strains are congruent, indicating co-speciation. (**B-D**) Mono-colonization rates, measured as CFU per polyp, for three *Curvibacter* strains across three different *Hydra* species after 7 days. (n=12). Statistical significance (ANOVA, followed by a Bonferroni post-test) is indicated by asterisks, *p < 0.05, **p < 0.01.

To test whether the investigated *Curvibacter* strains have co-adapted to their *Hydra* hosts we performed reciprocal recolonization experiments with three different *Hydra* species and their corresponding *Curvibacter* isolates (**Figure 1B-D**). All tested *Curvibacter* strains recolonized their native *Hydra* host best (**Figure 1B-D**). The highest differences in colonization rate were evident for the recolonization with *Curvibacter* AEP1.3. Interestingly, the isolates Hmag1.1 and Hvul1 establish similar colonization rates on *H. magnipapillata* and *H. vulgaris* (**Figure 1C, D**). This observation is consistent with the close relationship of both *Hydra* species (**Figure 1A**).

### Comparative genomics of free-living and *Hydra* associated *Curvibacter* strains

In the next step, we aimed at analyzing the co-speciation of *Curvibacter* to the *Hydra* host on the genomic level. For this purpose, we sequenced the genomes from two isolates of *Hydra* associated *Curvibacter* strains (Hmag1.1, Hvul1) in addition to the already available genome from the strain AEP1.3 (Pietschke et al., 2017). In public databases several free-living *Curvibacter* species are reported and their genomes are available (Ding & Yokota, 2004; Hahn et al., 2010; Lyu et al., 2024; Ma et al., 2016). Based on the genomes of three free-living strains (*Curvibacter delicatus* (GCA_041639495), *Curvibacter lanceolatus* (ATCC 14669) (GCF_000381265), *Curvibacter gracilis* (ATCC BAA-807) (GCF_000518645) and our three *Hydra*-associated strains (*Curvibacter* AEP1.3 (GCF_002163715), *Curvibacter* Hmag1.1, and *Curvibacter* Hvul1) a comparative genomic analysis was conducted (**Table S1**).

To assess genetic differences among all selected *Curvibacter* strains, an all-vs-all orthogroup (OG) overlap analysis was performed (**Figure 2A**). The three symbiotic *Curvibacter* strains exhibit the highest similarities to each other with 84 to 99% of common OGs. This pattern is also evident for the two free-living *Curvibacter* strains, *C. lanceolatus* and *C. gracilis*, which both possess 97% of common OGs. Both strains show less similarities when compared to the symbiotic species with 51 to 52% of shared OGs. The *C. delicatus* genome contains the fewest protein coding genes, resulting in the lowest number of OGs (**Table S2**). *C. lanceolatus* and *C. gracilis*, which have the highest number of OGs, share the fewest OGs with *C. delicatus*. In contrast, the OG distribution of *C. delicatus* exhibits the highest similarity compared to the other two free-living species, sharing 90% and 91% of its OGs, respectively. This result is potentially driven by the reduced protein-coding sequences of *C. delicatus*. In comparison, it shares 82 to 83% of its OGs with host-associated *Curvibacter* strains.

**Figure 2:**
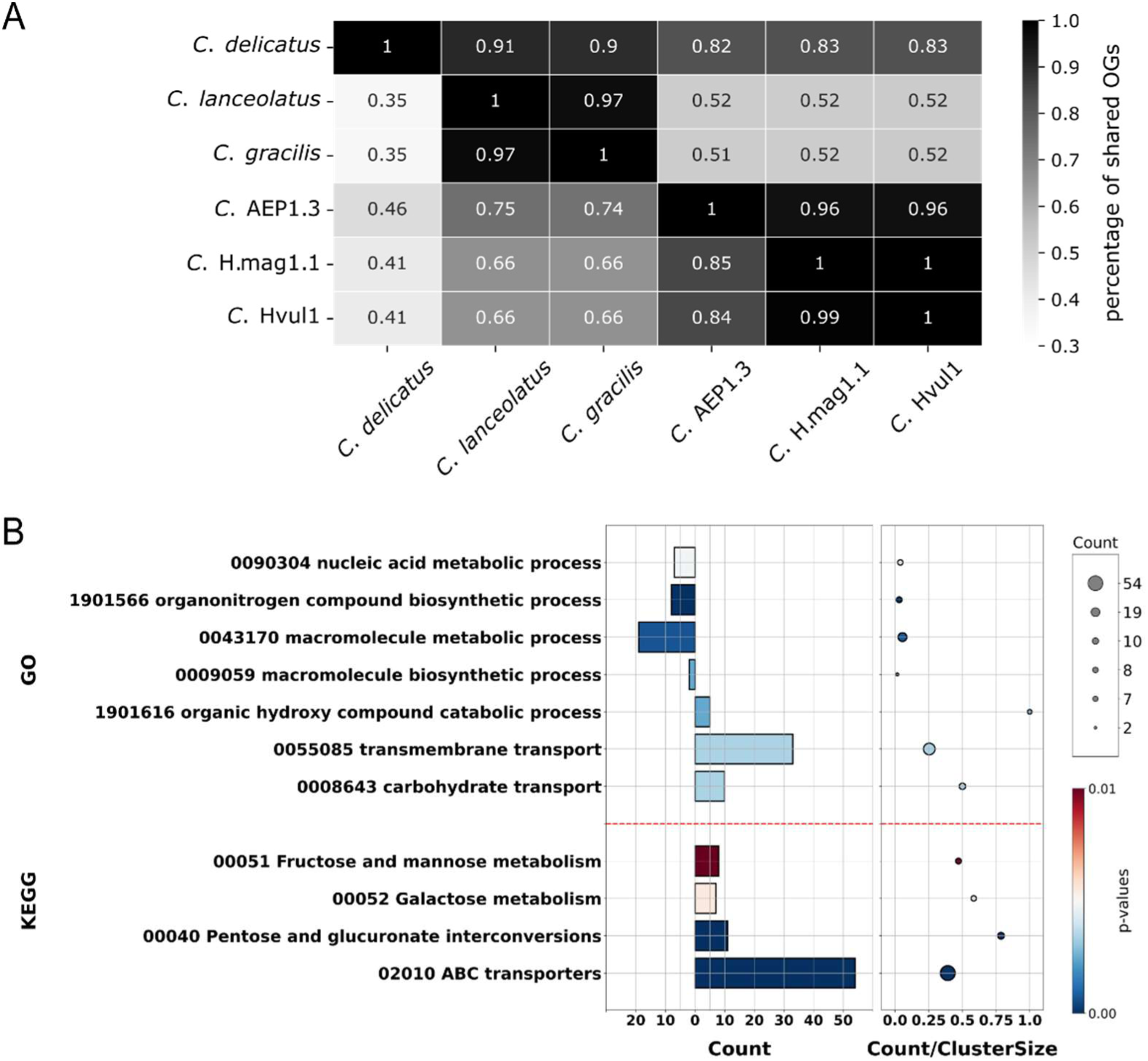
Ortholog and KEGG/GO analysis of *Curvibacter* proteomes. (**A**) Heatmap of orthogroup (OG) overlaps expressed as a percentage relative to the total OG count. The overlaps were normalized by dividing the OG overlap between two species by the total OG count of the species on the left axis. Highest similarities are observed within host-associated strains (*C.* AEP1.3, *C.* H.mag1.1 and *C.* Hvul1) and among the two free-living species *C. lanceolatus* and *C. gracilis*. (**B**) Selection of KEGG and GO enrichment analyses with the set of 693 protein sequences of *Curvibacter* AEP1.3 obtained from the symbiont specific set of OGs. The figure highlights a selection of the most notable categories, wherein the top four represent purified GO terms, and the remaining three correspond to enriched categories. The analyses reveal a significant enrichment of transporter-specific proteins (KEGG: ABC transporters; GO: carbohydrate transport and transmembrane transport). While three pathways related to sugar metabolism are enriched among the KEGG terms, the GO analysis highlights the enrichment of the organic hydroxy compound catabolic process, along with a purification of GO terms associated with the metabolic and biosynthetic pathways of macromolecules.

Subsequently, the OG data was filtered to identify OGs exclusive to the symbiotic strains. This analysis revealed 647 symbiont specific OGs (**Table S3**), with 693 of these genes present in *Curvibacter* AEP1.3. To assess functional enrichment within the symbiont specific OGs, a gene enrichment analysis was performed using the symbiont-specific protein sequence set of *Curvibacter* AEP1.3, incorporating both GO terms and KEGG Orthology (KO) (**Figure 2B**). Overall, four KEGG categories show significant enrichment, while 15 and 20 GO categories were identified as enriched and purified, respectively (**Table S4** and **S5**). Some GO terms form parent-child relationships within the GO hierarchy. Both enrichment analyses revealed a significant enrichment of transporter-specific proteins, particularly in the categories related to ABC transporters (KEGG), as well as carbohydrate and transmembrane transport (GO) (**Figure 2B**). Nearly 50% of transporters associated with carbohydrate transport (GO:0008643) are enriched, along with 45-75% of proteins involved in three KEGG pathways for sugar metabolism (ko00040, ko00052, and ko00051) (**Figure 2B**). Remarkably, all proteins within *Curvibacter* AEP1.3 associated with the organic hydroxy compound catabolic process category (GO:1901616) reside within the symbiont specific dataset. Additionally, proteins associated with macromolecule metabolism and biosynthesis (GO:0043170 and GO:0009059) exhibit significant purification. Taken together GO and KEGG terms associated with metabolic and transport pathways are significantly enriched and/or purified within the symbiont-specific gene set of 693 *Curvibacter* AEP1.3 genes, indicating that *Curvibacter* has undergone specific metabolic adaptations to thrive within its ecological niche in the glycocalyx of *Hydra*.

### Symbiont specific genes reveal associations with an extracellular polymeric substance (EPS) operon

Within the symbiont-specific OGs we identified one gene annotated as *epsH/exosortase B* (WP_087496557.1) among the purified GO category ‘macromolecule metabolic process’ that is part of an operon potentially linked to the biosynthesis and transport of extracellular polymeric substances (EPS) (Haft et al., 2006; Yoshida et al., 2003). The complete operon consists of eleven genes (**Figure 3A, Table S6**). Among those eleven genes, five genes reside in the symbiont*-*specific gene set (**Table S3,** EpsA, EpsL, EpsD, EpsH and EpsI).

**Figure 3:**
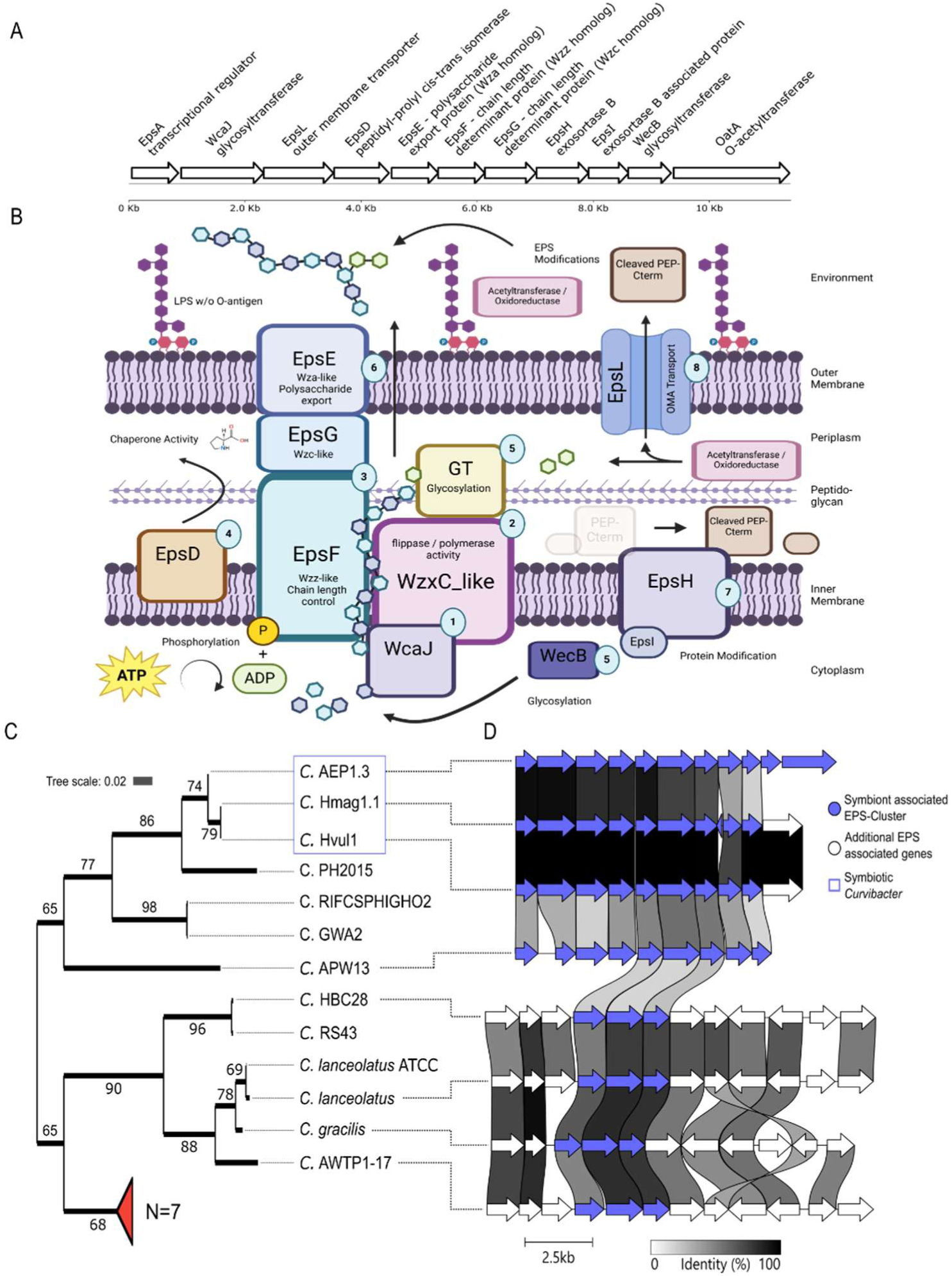
Putative functional roles of the EPS operon enzymes and a species-level phylogeny of *Curvibacter* reflecting the syntenic conservation of this operon. **(A)** Locus of the EPS operon of *Curvibacter* AEP1.3. **(B)** Putative functions of the EPS operon and neighboring genes in *Curvibacter* AEP1.3 suggest involvement in exopolysaccharide production and biofilm formation, with components such as EpsF, EpsG, EpsE, and WcaJ sharing homology with known Wzy-polymerase biosynthesis pathway proteins (Cuthbertson et al., 2009; Yoshida et al., 2003). **(1)** WcaJ may act as a priming glycosyltransferase and performs initial glycosylation (Pal et al., 2019; Patel et al., 2012). **(2)** In the Wzy-Polymerase biosynthesis pathway, the growing glycan is transported across the inner membrane into the periplasm by the Wzx flippase. There is no homologous protein within the EPS operon, but a neighboring enzyme, located nine genes downstream of the O-acetyltransferase of the EPS operon, has a WzxC domain (WP_087496544.1) and may act as a flippase. **(3)** The enzymes EpsF and EpsG share structural similarities with polysaccharide co-polymerase (PCP) proteins. EpsF contains a Wzz domain, while EpsG features a Wzc domain. The Wzz protein functions as a regulator of glycan chain length during polysaccharide synthesis, whereas the Wzc protein may modulates the overall activity and export readiness of the polysaccharide synthesis machinery through changes in its phosphorylation state (Kintz et al., 2008; Larue et al., 2009; Reid & Whitfield, 2005; Wugeditsch et al., 2001). **(4)** The EpsD enzyme is annotated as peptidyl-prolyl cis-trans isomerase, and it likely assists in the proper folding and stabilization of polysaccharide synthesis machinery proteins. **(5)** The EPS operon contains two glycosyltransferases, WcaJ and WecB, alongside four additional glycosyltransferases located in proximity (**Figure S1** and **Table S7)**. These enzymes are probably responsible for the sequential glycosylation of the polysaccharide, with each glycosyltransferase typically catalyzing the addition of a specific monosaccharide to the growing polymer chain, thereby contributing to the diversity and complexity of the polysaccharide structure. **(6)** Export of the polymer is likely mediated by the activity of EpsG and EpsE, which possesses a Wza domain. Wza acts as an outer membrane polysaccharide exporter, forming a channel for the translocation of the completed polysaccharide chain (Nesper et al., 2003). **(7)** The EpsH exosortase is proposed to cleave the N-terminal PEP-C-term motif of proteins. These proteins are typically associated with clusters of EPS synthesis (Haft et al., 2006). Along with other enzymes such as acetyltransferases or oxidoreductases, it is hypothesized that they act as EPS modifying enzymes. **(8)** EpsL is an outer membrane transporter with a beta barrel domain and a signal peptide. It is likely located at the outer membrane of *Curvibacter* AEP1.3 where it forms a pore for diffusion of other enzymes or molecules. The Wzy-polymerase itself is not present within the EPS operon, however, in the genome of *Curvibacter* AEP1.3 two genes possess Wzy_C_2 domains, the corresponding enzymes may act similar to Wzy. However, the exact polymerization process of the EPS produced by the operon remains unknown. The figure was created with https://BioRender.com. **(C)** Orthofinder species tree inferred by the STAG algorithm from sets of gene trees. The values at each bipartition represent the percentage of occurrences of that specific bipartition across the set of inferred species trees. The phylogeny is in direct comparison to a syntenic EPS region **(D)**. The synteny graph displays the conservation status of the eleven EPS operon genes of the three host-associated *Curvibacter* species and five representatives of the free-living *Curvibacter* species. The percentage identity of these genes decreases in free-living species. The red marked branch of the phylogeny shows a cluster of *Curvibacter* species without putative orthologous genes of the EPS operon.

The first gene in the operon, *epsA*, encodes a transcriptional regulator that contains an internal LuxR domain at its C-terminus but lacks an autoinducer-binding domain (**Figure 3A**). The second gene is homologous to *wcaJ*, a glycosyltransferase from *E. coli*, which functions as the priming glycosyltransferase in colanic acid polysaccharide synthesis (Pal et al., 2019; Patel et al., 2012). Many of these enzymes share similar domains with known components of the Wzy-polymerase biosynthetic machinery (Cuthbertson et al., 2009; Whitfield, 2006), including those responsible for regulation (EpsA), transport (EpsL and EpsE), chain length control (EpsF and EpsG), protein modification (EpsD and EpsH/EpsI), glycosylation (WcaJ and WecB), and acetylation (OatA) (**Figure 3B**).

To assess the presence of this potential EPS synthesis and transport operon within different *Curvibacter* strains in a higher resolution, we retrieved 14 additional assemblies from free-living *Curvibacter* strains (**Table S8**). Using the CATHI software tool (Becker et al., 2023), a reciprocal BLAST analysis of the eleven EPS operon protein sequences was conducted against a database consisting of the 20 *Curvibacter* protein coding genes. The analysis revealed that nine of the eleven protein sequences have reciprocal best hits (RBH) within all three *Hydra*-associated *Curvibacter* strains. Furthermore, only the minority of free-living species possessed RBHs to the eleven protein sequences (**Table S9)**.

Combining the OrthoFinder and synteny analysis we inferred a genome level phylogeny (Emms et al., 2018) (**Figure 3C**) as well as the conservation of the synteny of the EPS operon of all 20 *Curvibacter* strains (**Figure 3D**). All three *Hydra*-associated strains form a monophyletic cluster and exhibit a high synteny conservation of the EPS operon (**Figure 3C, D**). Together with four free-living strains the *Hydra*-associated strains form a sister group to all other *Curvibacter* strains (**Figure 3C**). Interestingly, these four free-living species, similar to the symbiotic species, show homology to nine genes within the operon. However, the percentage identity of corresponding enzymes is highly reduced compared to identities observed in *Hydra*-associated strains. The syntenic status of the EPS operon is reduced to three genes within six other free-living *Curvibacter*, while seven *Curvibacter* species within this cluster lack the entire operon (**Figure 3C, D**).

Taken together, most enzymes in the EPS operon, except for EpsE, EpsF and EpsG, lack syntenic conservation in free living *Curvibacter* strains. Furthermore, conserved proteins exhibit lower sequence identity in free-living species compared to symbiotic strains. These results suggest potential functional divergence driven by different evolutionary pressures in free-living and symbiotic environments. This conclusion is supported by the fact that many protein sequences that are syntenically conserved exclusively within the free-living Curvibacter strains (**Figure 3D**, white arrows) also shared homologies with enzymes associated with EPS production (**Table S10**). In addition, in symbiotic strains several neighboring genes are predicted to be involved in EPS production, primarily comprising glycosyltransferases, oxidases, and acetylases, which are essential for determining the final EPS structure (**Table S10**).

### *Curvibacter* AEP1.3 is producing extracellular polymeric substances

Based on the predicted functions (**Figure 3B**), it can be hypothesized that this operon is involved in the production of exopolysaccharide structures, such as capsular polysaccharides, glycosylated proteins and/or lipopolysaccharides (LPS). However, the exact functions of these genes remain unknown. Notably, two neighboring enzymes in the cluster are annotated as LPS biosynthesis protein (WP_087496544) and O-antigen ligase (WP_087496545). Consequently, an isolation of LPS was performed using the free-living *Curvibacter* strain JS11-12 and the symbiotic *Curvibacter* AEP1.3 (**Figure 4A**). The LPS was isolated from cultures grown in liquid R2A at 18°C, reaching the stationary phase. While *Curvibacter* AEP1.3 exhibits distinct bands corresponding to lipid A and core oligosaccharides in *E. coli* O55, bands corresponding to the long chain O-antigen are absent (**Figure 4A**). Interestingly, the LPS extraction of the free-living species, *Curvibacter* sp. JS11-12, shows distinct bands corresponding to the long chain O-antigen polysaccharide in *E. coli* (**Figure 4A**).

**Figure 4:**
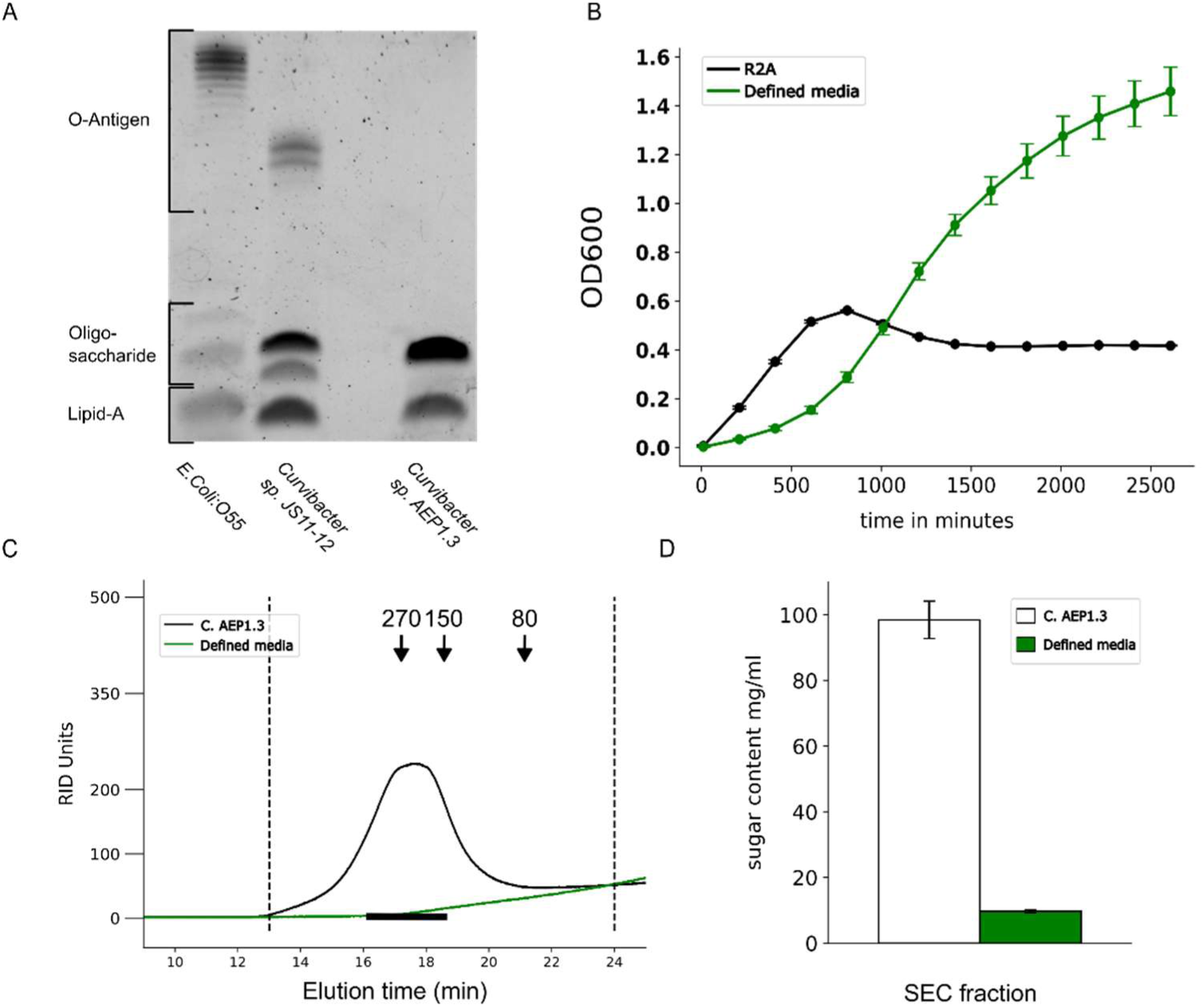
Analysis of EPS obtained from *Curvibacter* AEP1.3. **(A)** 15%-SDS gel electrophoresis of Hot-Phenol-Ether isolated lipopolysaccharide (LPS) fractions. *E. coli* LPS O55, provided by the gel staining kit, was used as a standard (left lane), while the two additional LPS samples were extracted from the free-living *Curvibacter* sp. JS11-12 strain (obtained from the DSMZ) and *Curvibacter* AEP1.3. The upper bands of *E. coli* O55 and *Curvibacter* sp. JS11-12 represent long chain O-antigen regions, corresponding to varying numbers of O-antigen repeat units. The lower bands indicate intermediate and short-chain O-antigen or oligosaccharides, while the lowest bands correspond to the lipid A core region of LPS (Jacobson et al., 2018; Wang et al., 2002). Separation of LPS from *Curvibacter* AEP1.3 does not result in visible bands for the long-chain O-antigen region. **(B)** Growth curve of *Curvibacter* AEP1.3 cultured in the standard bacterial-freshwater medium R2A (black curve) and in a defined media (green curve) at room temperature. Although the generation times are longer in the defined medium (**Figure S2**), *Curvibacter* AEP1.3 achieves higher OD values in this medium compared to R2A. **(C)** Size-Exclusion-Chromatographic analysis of the lyophilized and re-dissolved cell-free supernatant from *Curvibacter* AEP1.3 cultures (black), compared to lyophilized and re-dissolved defined medium as a negative control (green). The elution times of dextrose kilodalton standards are indicated by black arrows. In contrast to the negative control, both the Refractive Index Detector (RID) and the UV 280 nm detector (**Figure S3**) detected a signal in the higher molecular weight fractions of the SEC for the *Curvibacter* AEP1.3 wt supernatant. **(D)** Sugar content of *Curvibacter* AEP1.3 wt supernatant (white bars) and the defined media (green bars) as control of the higher molecular weight fractions obtained from the SEC measured with the Phenol-Sulfuric-Acid assay with glucose as standard (n=3). The detection of sugars in these fractions suggests that the high-molecular-weight molecules present in the supernatant are composed of polysaccharides.

Therefore, it is unlikely that the identified operon is responsible for the production of the O-antigen of LPS. In addition, the annotation and similarity of the potential EPS operon to the enzymes described by Yoshida et al. (2003) suggest the production of an exopolysaccharide that is not necessarily associated with the membrane. Therefore, we tested whether *Curvibacter* AEP1.3 secretes any sugar containing polymeric substances into the media. The growth medium R2A is not ideal for this purpose due to the presence of yeast extract and other polymeric substances, which influence the downstream analysis of the secreted substances. Therefore, we developed a defined growth medium for *Curvibacter* AEP1.3. Surprisingly, while exhibiting slightly reduced growth rates compared to R2A during logarithmic growth, the defined medium outcompetes R2A in terms of maximum OD_600_ values (**Figure 4B**).

In the next step, the defined medium was used to cultivate *Curvibacter* AEP1.3 at 18°C for 76 hours until the stationary phase was reached. Subsequently, all secreted polymeric substances were isolated through centrifugation, filtration, freeze-drying, and size-exclusion chromatography (SEC). SEC revealed two distinct peaks measured by the refractive index detector (RID). The first, smaller peak appeared at the onset of the exclusion at fraction 10 and extended to fraction 21 (**Figure 4C**). As this peak is not present in the media control (**Figure 4C**), it is likely that molecules eluting at this retention time correspond to high-molecular-weight molecules secreted specifically by *Curvibacter* AEP1.3. The second, broader and larger peak appeared also in the medium control and may correspond to monosaccharides and amino acids present in the medium. In addition, there were also two peaks detected with the UV detector in the *Curvibacter* AEP1.3 sample (**Figure S3**). The first peak, similar to the RID peak, indicates the presence of high-molecular-weight molecules that can absorb light at 280 nm, such as proteins or protein aggregates. The analysis of the carbohydrate concentration revealed a significantly higher sugar concentration in the *Curvibacter* AEP1.3 samples compared to the media control (**Figure 4D**). To summarize, while *Curvibacter* AEP1.3 has no distinct bands for long chain O-antigen polysaccharides, it produces and secretes polysaccharide based molecules into the medium.

### EPS gene knockouts impact growth behavior, monosaccharide composition, and recolonization efficiency

To investigate the role of the EPS operon in the interactions with *Hydra*, knockout mutants of *Curvibacter* AEP1.3 were generated for the *epsH* and *wcaJ* genes (**Figure 5A, B**). The epsH gene was selected because it occurs in the enrichment analysis, is present in the other symbiotic strains, and also has a very low number of RBHs in the reciprocal BLAST analysis. (**Figure S4** and **Table S9**). The *wcaJ* gene knockout was chosen due to its annotated function as a priming glycosyltransferase for the synthesis of the exopolysaccharide colanic acid (Pal et al., 2019; Patel et al., 2012).

**Figure 5:**
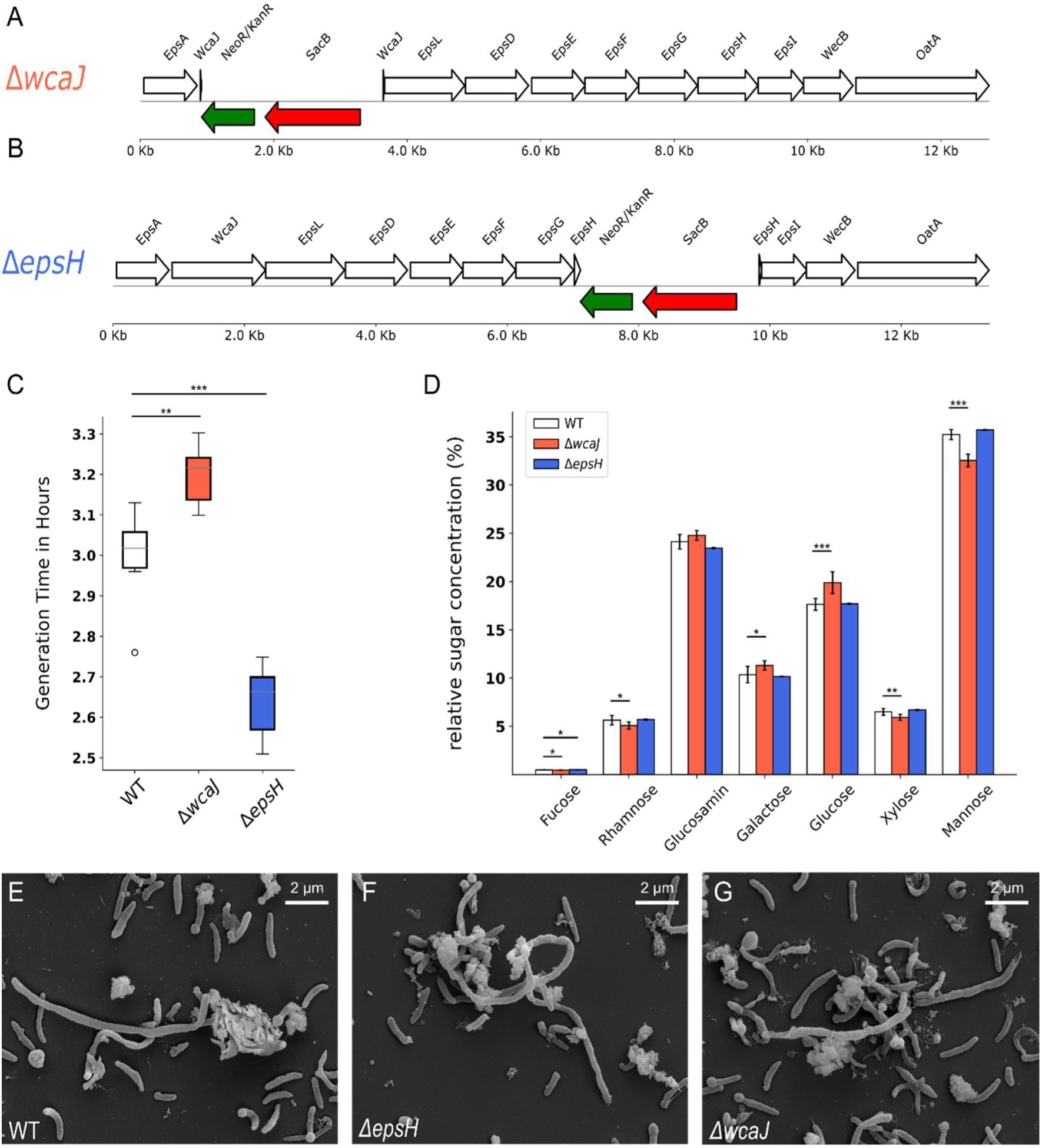
Comparison of morphology, growth and monosaccharide abundance of *Curvibacter* AEP1.3 wt and two mutant lines. (**A, B**) Visualization of gene knockouts within the EPS operon of *Curvibacter* AEP1.3 mutant lines *ΔwcaJ* (**A**) and *ΔepsH* (**B**). (**C**) Generation times for the wt, *ΔwcaJ*, and *ΔepsH* strains of *Curvibacter* AEP1.3 (n=6) were calculated using the logarithmic growth model during the exponential growth phase. Specifically, generation time in hour was assessed during the initial logarithmic growth phase, defined as the increase in optical density from OD_600_ 0.05 to OD_600_ 0.1. The mean generation time is significantly higher in *ΔepsH* mutant and significantly lower in the *ΔwcaJ* mutant compared to the wt strain. Statistical analyses were conducted using a one-way ANOVA, pairwise comparisons were conducted with Tukey’s HSD test to identify significant differences. Statistical significance is indicated by asterisks, with the following meanings: *p < 0.05, **p < 0.01, ***p < 0.001. The OD_600_ growth curve is provided within the supplementary material (**Figure S5**). (**D**) Relative monosaccharide abundance of isolated *ΔepsH*, *ΔwcaJ* and wt EPS. The abundances reflect the relative sugar concentrations obtained from both *ΔwcaJ* and wt cultures (n=7), along with *ΔepsH* (n=3). Significant differences can be observed in the Mannose and Glucose abundances between wt and *ΔwcaJ*. Statistical analyses were conducted using a two-way ANOVA, p-values were adjusted using the Bonferroni correction method. Statistical significance is indicated by asterisks, with the following meanings: *p < 0.05, **p < 0.01, ***p < 0.001. (**E - G**) Scanning-Electron-Microscopy (SEM) graph of *Curvibacter* AEP1.3 strains. There are no visible morphological differences among the *Curvibacter* AEP1.3 wt (**E**) and the mutant lines *ΔepsH* (**F**) and *ΔwcaJ* (**G**).

The *epsH* and *wcaJ* knockouts were generated using the pGT42 vector-mediated double crossover technique (Wein et al., 2018), resulting in mutant strains that contain a kanamycin resistance gene and a *sacB* gene inserted at the loci of the *epsH* and *wcaJ* genes (**Figure 5A, B**, green and red arrows, respectively). Interestingly, the knockout strains exhibited significant differences in growth behavior compared to the wt strain (**Figure 5C**). While the *ΔepsH* strain growth faster, the *ΔwcaJ* strain growth significantly slower compared to the wt-strain. To investigate potential alterations in EPS sugar composition in the *ΔepsH* and *ΔwcaJ* mutants, we subjected the content of the high molecular polymer peak isolated by SEC (**Figure 4C**, region within dashed lines) to HPAEC-PAD analysis, determining the EPS monosaccharide composition.

The analysis revealed significant differences in the relative sugar abundances of the *ΔwcaJ* mutant (**Figure 5D**) with a higher relative abundance of glucose and galactose and a lower relative amount of mannose, xylose and rhamnose compared to the wt.

In contrast, the *ΔepsH* mutant exhibited no significant changes in sugar composition, aside from a slight reduction in fucose levels. This finding suggests that, rather than directly affecting the synthesis and structure of EPS, EpsH may play a critical role in the proper assembly or secretion of surface proteins mediated by the selective cleavage of PEPC-term containing proteins. SEM (**Figure 5E-G**) revealed no visual morphological differences in the mutant strains compared to the wt strain.

To test the hypothesis that the EPS operon is involved in adaptation of *Curvibacter* to *Hydra,* mono-colonization experiments of germfree *Hydra* AEP were conducted (**Figure 6A**). The *ΔepsH* and *ΔwcaJ* mutants recolonized the germfree *Hydra* AEP polyps significantly worse compared to the wt strain (**Figure 6B**), supporting the hypothesis that the symbiont-specific EPS operon contributes to the co-speciation of *Hydra* and *Curvibacter*.

**Figure 6:**
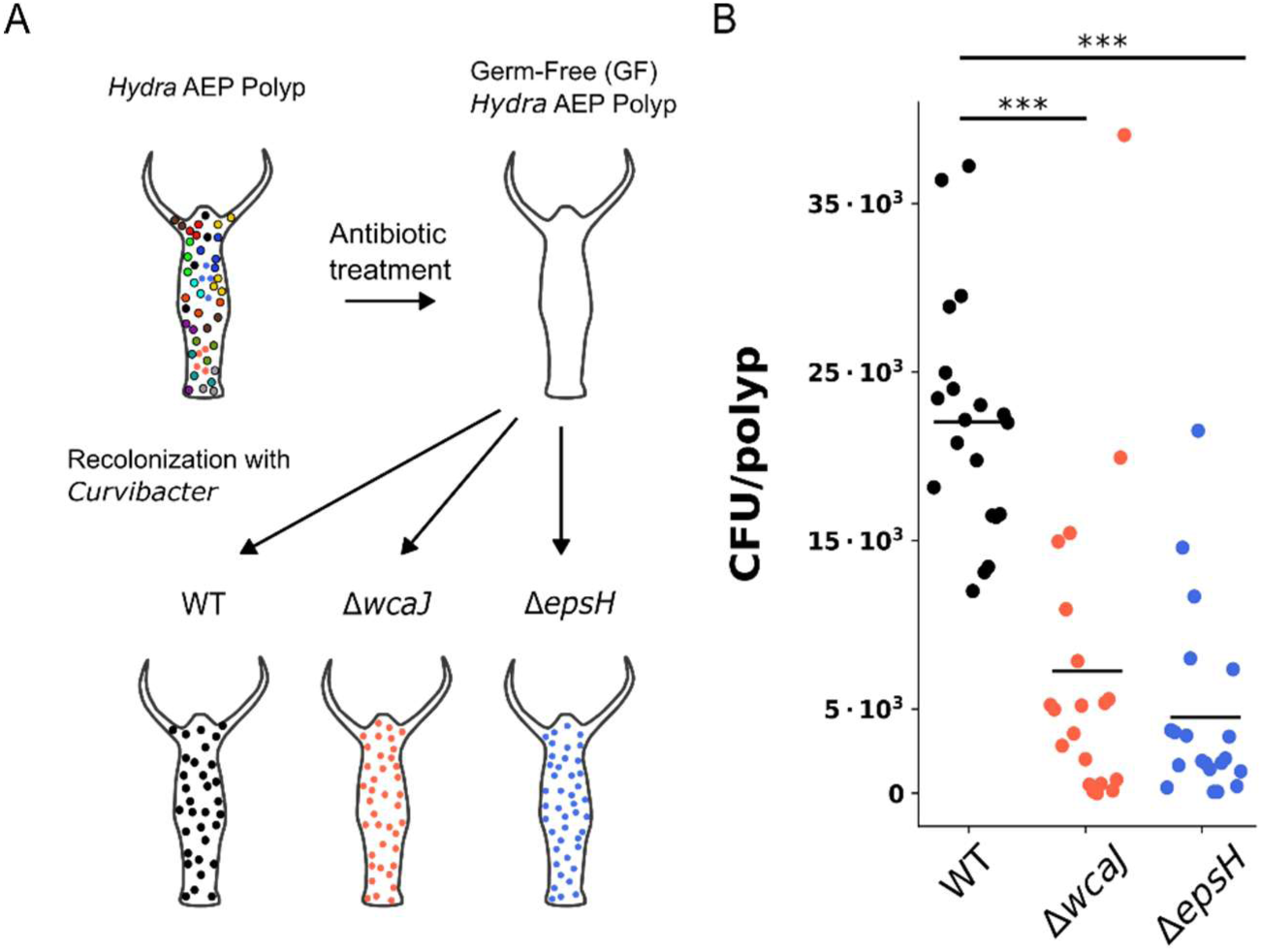
The recolonization efficiency of *Curvibacter* AEP1.3 mutant lines is impaired compared to the wt. **(A)** Scheme of recolonization of germ-free *Hydra* AEP polyps with *Curvibacter* AEP1.3 wt and the mutant strains *ΔwcaJ* and *ΔepsH*. **(B)** Recolonization rates after seven days recolonization, measured as CFU per polyp, for wt, *ΔwcaJ* and *ΔepsH* strains of *Curvibacter* AEP1.3. CFUs were counted for 20 polyps (n=20) per recolonization. The mutant strains exhibited significantly poorer performance compared to the wt strain; however, no significant differences were observed between the mutant lines themselves. Statistical analysis was performed using one-way ANOVA to compare group means, followed by pairwise t-tests with Bonferroni correction (***p < 0.001).

## Discussion

### Host-Symbiont Co-Speciation and Phylosymbiosis

The results presented in this study strongly support the concept of host-symbiont co-speciation between *Hydra* and its associated *Curvibacter* symbiont. Phylogenetic analyses revealed a congruent evolutionary pattern between the *Hydra* species and their bacterial symbionts of the genus *Curvibacter*, suggesting that evolutionary pressures have shaped the symbionts’ genomes, driving them towards mutual adaptation. This result aligns with observations obtained from the *Hydra viridissima* – *Chlorella* symbiosis, in which similar congruent phylogenies were reported (Kawaida et al., 2013). Native strains consistently outperformed non-native strains in recolonization experiments, emphasizing their specialized adaptations for colonizing their respective hosts. Since *Curvibacter* colonizes the surface of the ectodermal epithelium of *Hydra* polyps (Fraune et al., 2015), while it is in direct contact with the aquatic environment, it is of great interest to understand how the symbiosis between *Curvibacter* and *Hydra* is established and maintained.

Our findings shed light on broader implications for phylosymbiosis and coevolution in host-microbe systems (Moran & Sloan, 2015). Phylosymbiosis, the pattern where microbial community composition correlates with host phylogeny, has been observed across diverse host clades (Brooks et al., 2016). In mammals (Knowles et al., 2019), insects (Jackson et al., 2023; Qin et al., 2023), and invertebrates (O’brien et al., 2020; Pollock et al., 2018), intraspecific microbiome variation is consistently lower than interspecific variation, and microbial community similarities show significant topological congruence with host phylogenies (Brooks et al., 2016). Phylosymbiosis appears to be driven by host-microbiome interactions, as evidenced by reduced survival and performance in interspecific microbiota transplants (Brooks et al., 2016). While host selection of the microbiome is often cited as a mechanism, other ecological and evolutionary processes such as dispersal, drift, and diversification may also contribute to phylosymbiotic patterns (Kohl, 2020). In contrast to the community-level focus of phylosymbiosis, individual bacterial species often undergo coevolution with their hosts, resulting in highly specialized interactions. For instance, *Buchnera aphidicola* has coevolved with aphids, providing essential nutrients that the host cannot obtain from its diet (Douglas, 1998). This association, dating back 150 million years, has led to coevolution and codependence between the partners (Bennett & Moran, 2015). However, this obligate symbiosis also presents potential drawbacks, including metabolic costs, and symbiont genome degeneration, which may ultimately limit the ecological range and adaptability of aphid species (Bennett & Moran, 2015). Similarly, great ape species show complex co-evolutionary dynamics with individual bacteria. Several bacterial lineages in human and ape guts have co-evolved with their hosts over the last 15 million years, with divergence times consistent with hominid evolution (Moeller et al., 2016). These distinct examples of phylosymbiosis and individual bacterial coevolution highlight the complexity and diversity of evolutionary processes shaping host-microbe relationships, from broad community-level patterns to specific, finely tuned symbiotic interactions.

In the case of *Hydra*, the innate immune system’s ability to modulate microbial composition likely plays a pivotal role in maintaining phylosymbiosis (Franzenburg et al., 2013). By selectively fostering specific microbial communities by species-specific antimicrobial peptides, the host ensures compatibility and functional integration of its microbiota. On the other hand, individual bacterial symbionts, like *Curvibacter*, adapted to these host conditions ensuring the maintenance of association. By advancing our understanding of *Hydra*-symbiont co-speciation, this study contributes to the broader discourse on the evolutionary mechanisms underpinning host-microbiome dynamics. It raises compelling questions about how environmental pressures and host-specific traits influence microbial specialization, offering a framework for exploring the origins of phylosymbiosis within the *Hydra*-*Curvibacter* association.

### Bacterial Genomic Adaptations to Host Organisms

The comparative genomic analysis of symbiotic and free-living *Curvibacter* strains highlights key adaptations that enable *Curvibacter* to thrive within the *Hydra* host environment. Further evidence of these adaptations is the identification of orthogroups that are unique to symbiotic strains, many of which are associated with functional categories relevant to host colonization and survival. Hydra-associated strains exhibit a distinct enrichment of genes involved in carbohydrate transport. These adaptations likely support the acquisition and utilization of host-derived resources, enabling the symbionts to sustain their populations within the *Hydra* environment. Furthermore, genes involved in macromolecule biosynthesis and metabolic processes exhibit significant purification, reflecting distinct evolutionary pressures acting on a specific subset of these genes. This genomic specialization may underscore the importance of metabolic plasticity in facilitating the transition from free-living to host-associated lifestyles. It is also known from other host-bacteria models that host-associated microbes can undergo genetic diversification to exploit specific ecological niches. For instance, the coevolution of *Helicobacter pylori* with humans has resulted in lineage-specific adaptations that allow it to persist in the acidic environment of the stomach. The bacterium adapts to stomach acidity by increasing the use of positively charged amino acids in its membrane proteins (Xia & Palidwor, 2005). *H. pylori* exhibit niche-specific adaptations within the stomach allowing the colonization of approximately 50% of the human population (Ailloud et al., 2019), causing overt gastric disease in only a subset of infected individuals (Salama et al., 2013).

Similar to *Curvibacter*, *Lactobacillus reuteri* strains exhibit host-specific genomic traits that enhance their ability to colonize the gastrointestinal tracts of various mammals. Comparative genomic analyses have revealed broad genetic diversity among *L. reuteri* strains from different hosts, with distinct phylogenetic clades showing host specificity (Frese et al., 2011; J. Yu et al., 2018). Experimental studies in gnotobiotic mice have shown that *L. reuteri* has evolved host specialization, with rodent-associated strains possessing a specific genomic content that enhances their ecological performance in the mouse gut (Frese et al., 2011). The genomic events shaping host specificity also include the acquisition of genes encoding cell surface proteins, active carbohydrate enzymes, and other functional genes that contribute to adaptation to specific intestinal habitats (Frese et al., 2013; Wegmann et al., 2015; J. Yu et al., 2018). In summary, the genomic adaptations observed in *Curvibacter*, in addition to the known parallels to other host-associated bacteria, underline a central role in genetic specialization for symbiotic relationships and niche adaptation within the host environment.

### Role of Exopolysaccharides in Symbiont Adaptations

Exopolysaccharides play a critical role in mediating the specificity and stability of the *Hydra*-*Curvibacter* symbiosis. The identification of a conserved EPS operon in host-associated *Curvibacter* strains underscores its importance in facilitating microbial adhesion and colonization within the host’s glycocalyx. Functional analyses of this operon, particularly the *epsH* and *wcaJ* genes, revealed impaired recolonization capabilities of the corresponding mutant strains, suggesting that the production of specific glycans or glycoproteins is integral to maintaining the symbiotic relationship. These findings align with previous studies highlighting the role of EPS in microbial adhesion, biofilm formation (Vu et al., 2009), and immune evasion (Fanning et al., 2012; López et al., 2018; Watters et al., 2016) in other host-microbe systems (Gunn et al., 2016). EPS, primarily composed of polysaccharides and proteins form the matrix of microbial aggregates and biofilms (Sheng et al., 2010; Vu et al., 2009) mediate initial cell attachment to surfaces and protect bacteria from environmental stresses (Vu et al., 2009). In the context of host interactions EPS not only contribute to microbial adhesion and colonization but also play a crucial role in modulating host immune responses. Studies have shown that EPS can help symbiotic bacteria evade host immune detection by masking microbial-associated molecular patterns and inhibiting host immune receptors. For example, *Bifidobacterium breve* produces surface-associated EPS that enable evading adaptive B-cell responses and persistence in the gut (Fanning et al., 2012). Similarly, EPS from *Lactobacillus rhamnosus* forms a protective shield against host innate immune factors, enhancing its survival in the gastrointestinal tract (Lebeer et al., 2010). Furthermore, EPS produced by probiotic bacteria has been associated with various health benefits, including immune tolerance induction, anti-inflammatory effects, and protection against pathogens (Bhandary et al., 2023). Understanding the immunomodulatory properties of EPS in the *Hydra-Curvibacter* system could provide further insights into the evolution of host-microbe mutualisms and the molecular mechanisms underlying host immune tolerance.

EPS have also been shown to play pivotal roles in other host-symbiont interactions. In the squid-*Vibrio* symbiosis, for example, the EPS produced by the *syp* genes is essential for host colonization and biofilm formation (Shibata et al., 2012; Yip et al., 2005). Mutational studies have shown that most *syp* genes are required for successful colonization of the squid, with varying degrees of impact (Shibata et al., 2012). Similarly, in plant-rhizobia interactions, rhizobial EPS is essential for nodule formation and successful nitrogen fixation in leguminous plants (Acosta-Jurado et al., 2021). The plant receptor Epr3 can recognize and distinguish compatible and incompatible EPS in bacterial competition studies (Kawaharada et al., 2015). The EPS produced by *Lactobacillus plantarum* play a crucial role in strain-specific probiotic effects and host interactions. *L. plantarum* strains possess multiple gene clusters responsible for EPS production, with varying impacts on surface glycan composition (Remus et al., 2012). The removal of EPS can alter bacterial surface properties, stress survival, and immunomodulatory capacities in a strain-specific manner (Lee et al., 2016). The strain-specific nature of EPS contributes to the variability in probiotic efficacy observed among different strains (Bron et al., 2013).

Similarly, the conservation of EPS-related genes in symbiotic *Curvibacter* strains (**Figure 3C,D**) illustrates their adaptation to the *Hydra* niche. Comparative genomic analyses revealed that these genes are absent or poorly conserved in free-living strains, indicating their acquisition or specialization in host-associated strains. This specialization likely enhances the symbiont’s ability to persist in the host’s dynamic environment, which is characterized by antimicrobial peptides and other selective pressures. Moreover, the distinct sugar compositions of EPS from mutant strains, as well as the significant differences in generation times compared to the wt suggest that these polysaccharides play a specific role in mediating host-symbiont interactions, potentially influencing factors such as microbial adhesion and host immune modulation. Future studies should be aimed at determining the specific structure of the *Curvibacter* EPS polymer in more detail.

## Conclusion

This study elucidates the intricate dynamics of the *Hydra-Curvibacter* symbiosis, highlighting the evolutionary and molecular mechanisms that underpin this relationship. The evidence for host-symbiont co-speciation underscores the deep evolutionary connections between hosts and their symbionts, while the identification of specialized adaptations such as the EPS operon showcases the molecular innovations that enable *Curvibacter* to thrive in host environments. By integrating genomic and functional analyses, this research advances our understanding of the evolutionary processes that underpin host-microbe associations, offering valuable insights into the dynamics of microbial adaptation and the resilience of host-microbiome systems.

## Material & Methods

### Animal culture

Experiments were carried out using *Hydra vulgaris* (AEP) (hereafter *Hydra* AEP), *Hydra oligactis* (strain 10/02), *Hydra viridissima* (strain A99), *Hydra magnipapillata* (strain 105), *Hydra vulgaris* (strain Basel), and *Hydra circumcincta* (strain M7). All animals were cultured under constant, identical environmental conditions including culture medium, food (first-instar larvae of *Artemia salina*, fed three times per week) and temperature according to standard procedures (Lenhoff & Brown, 1970). For all experiments, adult polyps without buds or gonads were used.

### Isolation of *Curvibacter* strains

Single *Hydra* polyps from each species were placed in a 1.5-ml reaction tube and washed three times with 1 ml sterile filtered *Hydra* medium. After homogenization with a pestle, 100 μl (equates to 1/10 of a polyp) was plated on Reasoner’s 2A (R2A) agar plates (Sigma-Aldrich). After incubation at 18 °C for 5 days, single colony-forming units (CFUs) were isolated and cultivated in liquid R2A medium. The bacteria were identified by Sanger sequencing of the 16S rRNA gene using the universal primers Eub-27F and Eub-1492R (Weisburg et al., 1991) and stocks were stored in Roti-Store cryo vials (Carl Roth, Karlsruhe, Germany) at −80 °C.

### Phylogenetic

For Hydra, COI sequences of the desired lineages were acquired from the NCBI database: *H. vulgaris* (AEP) (EF059935), *H. carnea* (EF059940), *H. magnipapillata* (EF059934), *H. vulgaris* (EF059936), *H. oligactis* (EF059937), *H. circumcincta* (EF059938), and*H. viridissima* (EF059941). Sequence alignment for the cytochrome oxidase genes was generated using Clustal W incorporated in MEGA11 sequence analysis software package (Tamura et al., 2021). A model test was used to estimate the best-fit substitution models for phylogenetic analyses. For the maximum-likelihood analyses, genes were tested using the General Time Reversible (GTR + I) model. A bootstrap test with 1,000 replicates for maximum likelihood using a random seed was conducted.

For *Curvibacter*, sequences for 16S rDNA were acquired of all isolates via Sanger-sequencing using the universal primers Eub-27F and Eub-1492R (Weisburg et al., 1991). Evolutionary analysis was conducted in MEGA11 (Tamura et al., 2021). All sequences were aligned using the integrated Clustal W alignment option with default parameters. The evolutionary history was inferred by using the Maximum Likelihood method and Hasegawa-Kishino-Yano model (Hasegawa et al., 1985). A discrete Gamma distribution was used to model evolutionary rate differences among sites. Bootstrap values were calculated based on 100 replicates.

### Generation of germfree *Hydra*

Polyps were transferred to a sterile beaker containing 30 ml of sterile S-medium supplemented with antibiotic solutions (50 µg/ml each of ampicillin, rifampicin, spectinomycin, streptomycin, and neomycin). To assess any potential impact of DMSO, which was used as a solvent for rifampicin, control polyps were treated with DMSO (1 µl/ml). The beaker was sealed airtight and stored at 18°C in the dark. Over ten days, the medium and antibiotic solution were replaced every two days under sterile conditions as previously described (Franzenburg et al., 2012). Following antibiotic treatment, the GF polyps were washed with sterile S-medium, transferred to a fresh beaker, and incubated for an additional two days. As a contamination control, two polyps from each batch were homogenized in 100 µl of sterile S-medium using zirconia beads in a screw-cap tube and plated on R2A-Agar (ROTH) plates. After 3 days of incubation at room temperature, absence of CFUs indicated successful antibiotic treatment (Franzenburg et al., 2012). For culture-independent analysis, total DNA was extracted from single polyps using the DNeasy Blood & Tissue Kit (Qiagen). The 16S rRNA genes were amplified using the universal primers Eub-27F and Eub-1492R (Weisburg et al., 1991) in a 30-cycle PCR. Sterility was verified by the absence of a PCR-product.

### Recolonization experiments

*Curvibacter* isolates were cultured in liquid R2A medium for 3 days at 18 °C. Following centrifugation at 1380 × *g* for 10 min, the bacterial pellet was resuspended in sterile *Hydra* medium and optical density (OD_600_) was measured. For mono-association experiments, polyps were recolonized in *Hydra*-Medium with 5,000 *Curvibacter* cells per ml. For the recolonization experiments with the *Curvibacter* AEP1.3 mutants, polyps were recolonized with 50,000 *Curvibacter* cells per ml. Non-associated bacteria were removed by washing with sterile *Hydra* medium after 24 hours. After 7 days of recolonization, the S-medium was removed, and the *Hydra* polyps were carefully washed with sterile S-medium to eliminate residual bacteria. Each polyp was then transferred to an individual 1.5 ml screw-cap tube containing 100 µl of sterile S-medium. Subsequently, zirconia beads (1 mm) were added to the tubes, and the polyps were homogenized using a shaking homogenizer (BeadBug^TM^). Homogenates were serially diluted (1:20 and 1:40), and 100 µl of each dilution was plated onto R2A-Agar plates. The plates were incubated at room temperature for 72 hours, after which colony-forming units (CFUs) were counted and adjusted for the respective dilution factors. For the mono-association experiments, statistical analysis of the bacterial load was conducted using one-way analysis of variance (ANOVA). Dunnett’s test was used as a *post hoc* test to compare treatment with control samples. For the recolonization experiments with the *Curvibacter* AEP1.3 mutants, statistical analysis was performed using one-way ANOVA to compare group means, followed by pairwise t-tests with Bonferroni correction.

### Genome sequencing

For sequencing the genomes of the two *Curvibacter* isolates *Curvibacter* Hmag1.1 and *Curvibacter* Hvul1 paired-end libraries were prepared using an Illumina TruSeq LT kit with a median fragment size of 402 bp. Mate-pair libraries were prepared using an Illumina Nextera mate-pair kit with an insert size of 7.3 kb. The libraries were sequenced on a MiSeq instrument at the Biomolecular Resource Facility, The Australian National University, Canberra, Australia. The reads were quality trimmed and adaptors were clipped using libngs (https://github.com/sylvainforet/libngs). Mate-pair libraries were processed using NextClip (Leggett et al., 2014), keeping only read pairs in which the Nextera adaptor was found. The processed reads were assembled with SPAdes v3.5.0 (Bankevich et al., 2012). Gaps were filled using GapFiller v1-11 (Nadalin et al., 2012). Genes were predicted using Prokka (Seemann, 2014). Genome sequences were uploaded to NCBI as nucleotide FASTA files and are deposited under the BioProject identifier PRJNA1232435.

### Comparative genomic analysis

The E-Direct (22.1) software from NCBI was used to search and retrieve information on available *Curvibacter* assemblies. In total there were 48 assemblies. These assemblies were then filtered by the assembly completeness level, all assemblies annotated as “Contigs” were removed, resulting in a set of 18 *Curvibacter* assemblies. The *Curvibacter* assemblies (Supplementary Table S1, S8) were downloaded from NCBI. The downloaded nucleotide FASTA files were used as input for Prokka (1.14.6) to generate GenBank and proteome FASTA files (Seemann, 2014). The resulting proteomes together with the two proteome FASTA files of *Curvibacter* Hmag1.1 and *Curvibacter* Hvul1 were used as input for CATHI (Becker et al., 2023) and OrthoFinder (2.5.5) (Emms & Kelly, 2015, 2019). The synteny analysis was performed using clinker (0.0.27) (Gilchrist & Chooi, 2021). Clinker requires GenBank files that exclusively contain the gene regions of interest. To meet this requirement, we developed a custom Python script designed to extract relevant gene regions from the GenBank files generated by Prokka. The script identifies these regions based on reciprocal best hits (RBHs) obtained through the CATHI pipeline, particularly we checked the synteny status of the genetic loci around RBHs of EpsE (WP_087496560.1), EpsF (WP_087496559.1) and EpsG (WP_087496558.1). Gene sequences within the synteny, that do not share identities above 25% with *Curvibacter* AEP1.3 were subjected to an additional BLAST analysis. This BLAST analysis was conducted using CATHI with a subset of the Reference Proteins database, containing only high-quality bacterial proteomes (assembly status “Complete Genome” or “Chromosome”), and an e-value cut-off of 0.05. The species tree was generated based on a set of orthologous gene trees by OrthoFinder, which is using the STAG algorithm (Emms et al., 2018). OrthoFinder results were further used for an orthogroup (OG) overlap analysis, to check the evolutionary conservatism among the tested *Curvibacter* strains. To minimize bias arising from differences in total OG count, the number of shared OGs between each pair of strains was normalized by dividing it with the total number of OGs in each strain, reciprocally.

The inference of the symbiont specific OG set was done using a custom Python script using the OG table provided by OrthoFinder. Kyoto Encyclopedia of Genes and Genomes (KEGG) annotations were retrieved using the BlastKOALA (Kanehisa et al., 2016) webservice with the *Curvibacter* AEP1.3 proteome as input file. Gene Ontology (GO) terms were retrieved by parsing the public available Reference Sequence database GFF annotation file of *Curvibacter* AEP1.3 (GCF_002163715.1). The KEGG enrichment analysis was performed using the clusterProfiler package (3.20) (G. Yu et al., 2012) of R, the GO analysis was performed using the goatools (Klopfenstein et al., 2018) Python package.

### Preparation of defined medium

The defined medium was formulated using the M9 recipe as a template, with salt concentrations adjusted to match those of R2A. To support *Curvibacter* growth, nine amino acids and ammonium were added. The medium was buffered with HEPES to counteract acidification caused by *Curvibacter* metabolism. The M9 salt, trace elements, biotin, thiamine, NH₄Cl, HEPES, MgSO₄, CaCl₂ solutions, along with distilled water, were autoclaved. The 40% glucose and L-amino acid solutions were filter-sterilized using a 0.22 µm filter. All chemicals were stored at 4 °C prior to use. The resulting solution was sterilized by filtration using a 0.22 µm bottle-top filter to remove any potential microbial contaminants.

**Table 1:** Components of the defined medium.

| Chemical | Concentration in media |
| --- | --- |
| Na<sub>2</sub>HPO<sub>4</sub> | 337 µM |
| KH<sub>2</sub>PO<sub>4</sub> | 220 µM |
| NaCl | 855 µM |
| NH<sub>4</sub>Cl | 935 µM |
| Glucose | ~ 22,2 mM |
| CaCl<sub>2</sub> | 300 µM |
| MgSO<sub>4</sub> | 1 mM |
| HEPES (C<sub>8</sub>H<sub>18</sub>N<sub>2</sub>O<sub>4</sub>S) | 10 mM |
| NH<sub>4</sub>Cl | 10 mM |
| Histidine | 100 mg/L |
| Methionine | 100 mg/L |
| Asparagine | 100 mg/L |
| Arginine | 100 mg/L |
| Tryptophane | 100 mg/L |
| Isoleucine | 100 mg/L |
| Phenylalanine | 100 mg/L |
| Threonine | 100 mg/L |
| Aspartate | 100 mg/L |
| Biotin | 1 mg/L |
| Thiamin | 1 mg/L |
| EDTA | 134 µM |
| FeCl<sub>3</sub>·6H<sub>2</sub>O | 31 µM |
| ZnCl<sub>2</sub> | 6,2 µM |
| CuCl<sub>2</sub>·2H<sub>2</sub>O | 0,76 µM |
| CoCl<sub>2</sub>·2H<sub>2</sub>O | 0,42 µM |
| H<sub>3</sub>BO<sub>3</sub> | 1,62 µM |
| MnCl<sub>2</sub>·4H<sub>2</sub>O | 0,081 µM |
| H<sub>2</sub>O | - |

### Generation of *Curvibacter* AEP1.3 knockout strains

Pre-cultures of *Curvibacter* AEP1.3 were grown in either liquid R2A medium or defined medium at 18°C at 170 RPM. *Curvibacter* AEP1.3 cultures in R2A medium were grown for approximately two days, while cultures in defined medium were cultivated for around 3 days, before entering the stationary phase. *Escherichia coli (E. coli)* DH5α and *E. coli* DH5α WM3064 (*E. coli* ΔDAP) strains were revived from cryo-cultures onto LB-Agar plates. For the *E. coli* ΔDAP strain, 0.1 mM Diaminopimelic acid (DAP) was additionally included to support growth. For plasmid isolation, conjugation, and knockout experiments, *E. coli* cultures were inoculated from LB-Agar plates into liquid LB medium. The medium was supplemented with the appropriate antibiotics and/or DAP. *E. coli* cultures grew overnight at 37°C with continuous shaking at 170 RPM.

The deletion of the *epsH* and *wcaJ* gene regions was performed using a homologous recombination strategy as described in (Wein et al., 2018). To construct the desired plasmid, the first flanking region was introduced into the vector pGT42. The insertion was achieved through blunt-end ligation following restriction digestion with ScaI, designed to restore the stop codon of the ampicillin resistance gene. The resulting plasmid solution was transformed into chemically competent *E. coli* DH5α using the heat-shock method. Transformants were screened on LB-agar plates supplemented with 500 µg/ml ampicillin and 25 µg/ml chloramphenicol. Colony PCR was performed to confirm successful integration of the first flanking region. Positive clones were cultured overnight in liquid LB medium supplemented with ampicillin and chloramphenicol. Plasmid DNA was isolated using the NucleoSpin Plasmid Quick Pure Kit (Macherey-Nagel, Germany) and verified by sequencing. Next, the second flanking region was introduced into the validated pGT42 plasmid via a similar restriction-ligation process using HpaI. Cultures were treated with kanamycin for selection, and the resulting plasmid was confirmed via colony PCR, plasmid isolation, and sequencing as described above. The validated pGT42 vectors, containing both flanking regions, were introduced into *E. coli* ΔDAP cells using the heat-shock method. Transformants were screened via colony PCR, and positive clones were subsequently used for biparental mating with *Curvibacter* AEP1.3. The *epsH* and *wcaJ* gene targeting vectors are provided as GenBank files in the GitHub repository of this project. For conjugation, *Curvibacter* AEP1.3 (5 ml) and the *E. coli* ΔDAP donor strain (3 ml) were grown in liquid R2A medium to stationary phase. The *E. coli* ΔDAP culture was centrifuged at 3,000 g for 3 minutes, washed with liquid R2A, and centrifuged again. The *Curvibacter* AEP1.3 culture was then added, and the combined suspension was centrifuged at 3,000 g for 3 minutes. The pellet was resuspended in 700 µl R2A supplemented with 0.1 mM DAP and incubated for 3–6 hours at 30°C without shaking. A 100 µl aliquot of the conjugation suspension was spotted onto R2A-Agar plates with 0.1 mM DAP and incubated for 16–24 hours. Cells were then scraped from the plate, resuspended in 600 µl R2A (without DAP), and 100 µl of the suspension was spread on R2A-Agar containing 5 µg/ml kanamycin. After 48 hours, *Curvibacter* AEP1.3 clones were isolated and screened for integration of the kanamycin and SacB cassettes from pGT42, as well as for successful removal of the targeted gene regions by colony PCR.

### CLARIOstar growth experiments

Growth experiments were performed using the CLARIOstar plate reader (BMG Labtech). Pre-cultures for growth assays were diluted to an OD_600_ of 0.05 in 800 µl defined medium per well and inoculated into 48-well plates. The plate reader was programmed to measure OD_600_ values every 10 minutes throughout the experiment. For each growth experiment, at least three wells were inoculated with 800 µl of defined medium alone to serve as controls for contamination assessment and to measure blank values. The experiments were conducted at room temperature (∼21°C – 24°C – due to shaking) with continuous shaking of 500 RPM between measurements. The generation time g was calculated based on the standard exponential bacterial growth model, assuming bifurcations.

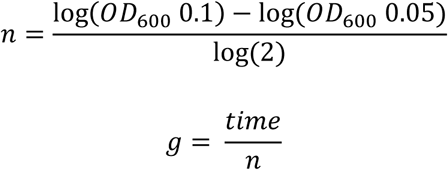

The number of generations during the time of bacterial growth is denoted by the number n, time of bacterial growth is based on the duration of how long the bacteria grow from an OD_600_ 0.05 to 0.1.

Statistical analysis included the Shapiro test to assess normal distribution, the Levene test to evaluate variance differences, and one-way ANOVA to determine differences in mean values. Pairwise comparisons were conducted using Tukey’s HSD (honestly significant difference) test.

### Scanning Electron Microscopy

*Curvibacter* AEP1.3 wt and the mutant strains *ΔepsH* and *ΔwcaJ* were grown in liquid defined medium at 18°C for 72 hours. Before Scanning Electron Microscopy (SEM) sample preparation, bacterial cultures were diluted to an OD_600_ of 1.6. For bacterial cell fixation, cover glasses (1 cm diameter) were coated with Poly-L-Lysine. Prior to coating, the cover glasses were washed with 70% ethanol and dried using compressed air. Cover glasses were then placed into 24-well plates and incubated for 5 minutes in a 0.1% (w/v) Poly-L-Lysine solution prepared in distilled water. The coated cover glasses were dried with compressed air and transferred to a new 24-well plate with the coated side facing upwards. Each well was filled with 1 ml of defined medium. Subsequently, 20 µL of bacterial cultures or defined medium (as a control) was added to the wells containing the coated cover glasses. The solutions were gently mixed by pipetting. The 24-well plate was then centrifuged at 1,500 g for 15 minutes at 4°C.

The medium was removed, and samples were incubated for 1 hour in 1 mL of fixation solution (0.1 M sodium cacodylate buffer, 2.5% glutaraldehyde, 2% formaldehyde in distilled water). Samples were then washed four times with washing solution (0.1 M sodium cacodylate buffer in distilled water). To ensure lipid fixation, an additional step was performed using 1 mL of osmium tetroxide fixation solution (1% OsO₄, 0.1 M sodium cacodylate buffer in distilled water) per sample, incubating for 1 hour at 4°C in the dark. Afterward, samples were washed four more times with washing solution and dehydrated through a graded ethanol series (30%, 50%, 70%, 80%, 90%, 96%, and 100%). Each ethanol solution was applied twice for 15 minutes, starting with 30%. After dehydration, samples were immersed in 100% ethanol and subjected to critical point drying. Therefore, samples were washed six times with liquid CO_2_ and the remaining CO_2_ was slowly evaporated in a pressurized chamber to remove all traces of liquid EtOH.

Next, samples were transferred to a sample holder with an adhesive pad using tweezers. The samples were subjected to sputter coating, during which a gold layer was deposited using plasma at a pressure of 0.08–0.09 mbar in an argon atmosphere. Finally, the gold coated samples were transferred to the SEM Zeiss SUPRA 55VP machine for imaging.

### Isolation, separation and staining of bacterial lipopolysaccharides

Lipopolysaccharides (LPS) from *Curvibacter* sp. JS11-12 and *Curvibacter* AEP1.3 were isolated and separated using the method described in (Davis & Goldberg, 2012). The bacterial cultures were incubated for 72 hours. The culture was diluted to OD_600_ of 0.5 and an aliquot of 1.5 ml was centrifuged at 10,600 g for 10 minutes and the pelleted bacteria were subjected to the LPS extraction protocol. Bacterial pellets were resuspended in 200 μl of 1×SDS lysis buffer (2% β-mercaptoethanol (BME), 2% SDS, 10% glycerol in 50 mM Tris-HCl (pH 6.8) and bromophenol blue dye) by gentle pipetting to ensure complete resuspension. The samples were boiled in a water bath at 100°C for 15 minutes and cooled to room temperature. To each sample, 5 μl each of DNase I and RNase solutions (238 µg/ml each) were added, followed by incubation at 37°C for 30 minutes. Proteinase K solution (455 µg/ml) was then added and incubated at 59°C overnight. Ice-cold Tris-saturated phenol (200 μl) was added to each sample. Tubes were tightly capped and vortexed for 5–10 seconds. The samples were incubated at 65°C for 15 minutes with occasional vortexing, cooled to room temperature, and mixed with 1 ml of room-temperature diethyl ether. The samples were vortexed for 5–10 seconds and centrifuged at 20,600 g for 10 minutes. The bottom blue layer was carefully extracted, avoiding contamination from the upper clear layer. A small amount of the blue layer was intentionally left behind to minimize contamination risk. This extraction process was repeated two times. If samples remained cloudy, additional extractions were performed as needed. Following extraction, 200 μl of 2×SDS lysis buffer was added to each sample and stored at 4°C. Samples were separated on 15% SDS-polyacrylamide gel, using 12 μl of prepared LPS per lane for visualization.

LPS fractions were stained using the Pro-Q™ Emerald 300 Lipopolysaccharide Gel Stain Kit (Thermo Fisher Scientific). The *E. coli* O55 LPS standard (provided within the Gel Stain Kit) was diluted 10-fold (to 250 µg/ml) in 2×SDS. The 15% SDS-polyacrylamide gel was immersed in a fixation solution (50% methanol and 5% acetic acid in distilled water) with gentle agitation on an orbital shaker for 45 minutes to fixate LPS. The fixation solution was carefully removed, and the process was repeated. The gel was incubated in the wash solution (3% glacial acetic acid in distilled water) for 15 minutes with gentle agitation on an orbital shaker. The washing solution was then removed, and the step was repeated. To oxidize LPS carbohydrates, the gel was immersed in the oxidizing solution (periodic acid as provided in the Kit and 250 ml of 3% acetic acid) by gentle agitation for 30 minutes. Following oxidation, the gel was washed twice as described earlier. A staining buffer was prepared by adding 500 µL of Pro-Q® Emerald 300 stock solution to 25 ml of staining solution. The buffer was applied to the gel and incubated for 120 minutes with gentle agitation in the dark. The gel was then washed twice with the washing solution, following the previously described procedure. The LPS fractions were visualized using the Image Lab (Bio-Rad) software and the ChemiDoc XRS+ (Bio-Rad) imaging system.

### EPS extraction

The growth of *Curvibacter* AEP1.3 for EPS extraction for carbohydrate analysis were carried out in a 50 ml culture in Erlenmeyer flasks at 18°C, with continuous shaking at 170 RPM in a New Brunswick Innova 42 incubator for 72 hours. For the monosaccharide analysis, *Curvibacter* AEP1.3 growth for EPS extraction was conducted using the Multi-Cultivator MC-1000-OD (PSI). Each vial was inoculated with 60 ml of *Curvibacter* AEP1.3 pre-culture at an OD_600_ of 0.1. For each experiment, at least one vial was filled with defined medium as a control. Growth in the MC device was carried out in the dark at 18°C. Cultures grew for nine days to ensure higher concentrations of EPS. After growth, each sample was divided into two 50 ml falcon tubes and centrifuged at 4,500 g for 10 minutes. Next, the supernatant was filtered through a 0.22 µm filter and transferred to a fresh 50 ml falcon. The samples were stored at - 20°C until further use. The cell-free supernatant was freeze-dried. Each lyophilized sample was then redissolved in 700 µl of autoclaved, distilled H₂O before combining pairs of samples. The resulting 1.4 mL solution was subjected to size exclusion chromatography (SEC) by applying the sample to a Superose 12 10/300 GL (Cytiva) column, separating small molecules from the defined medium and bacterial macromolecules. Fractions containing macromolecular components were pooled and vacuum-dried for carbohydrate analyses.

### Analysis of total sugar abundance

Vacuum-dried samples were redissolved in 1 ml of autoclaved distilled water. Total sugar abundance was quantified using the Phenol-Sulfuric Acid method, as described by (Masuko et al., 2005) using a glucose standard curve. In a 96-well plate, 50 µl of each sample or standard was added in triplicate. To each well, 150 µl of concentrated sulfuric acid was added, and the solution was mixed by pipetting. Subsequently, 30 µl of a 5% phenol solution was added to each well, and the mixture was again pipetted to ensure homogeneity. The plate was incubated at 90°C for 10 minutes to facilitate the reaction and then cooled to room temperature for approximately 30 minutes. Absorbance values for the samples and the standard series were measured across a wavelength range of 400–550 nm using a CLARIOstar plate reader to determine total sugar concentrations. Glucose concentrations in the samples were calculated based on the absorbance at 490 nm.

### Monosaccharide analysis

Monosaccharide composition analysis was conducted using high-performance anion-exchange chromatography with pulsed amperometric detection (HPAEC-PAD). Pooled macromolecular fractions containing the EPS were subjected to trifluoroacetic acid (TFA) hydrolysis. For the hydrolysis the samples were dissolved in 2 M aqueous TFA and heated at 121°C for 90 minutes, after which they were cooled on ice and centrifuged at 12,000 RPM for 5 minutes. The remaining acid was evaporated under a constant airflow at 40°C for approximately 80 minutes. Following this, 300 µl of isopropanol was added, the solutions were vortexed and dried again under a constant airflow at 40°C for 15 minutes. This step was repeated a second time. The samples were then dissolved in 500 µl water and subjected to anion-exchange chromatography utilizing a Knauer Azura HPAEC system (Knauer, Berlin). The CarboPac PA20 3×30 mm column was used as pre-column, while the CarboPac PA20 3×150 mm column was used as main column (Thermo Fisher). As a flow rate 0,4 ml/min was used. The gradient profile for eluents is described in the following table.

**Table 2:** Gradient profile of the HPAEC-PAD analysis.

| Time in minutes | 2 mM NaOH in H<sub>2</sub>O as % | 150 mM NaOH in H<sub>2</sub>O as % |
| --- | --- | --- |
| Initial | 100 | 0 |
| 21 | 100 | 0 |
| 22 | 0 | 100 |
| 25 | 0 | 100 |
| 26 | 100 | 0 |
| 35 | 100 | 0 |

Data acquisition and integration were performed using the ClarityChrom software (7.4.2.107, Knauer). The obtained data was subjected to a two-way ANOVA statistical test followed by a Bonferroni post-test.

### Computational scripts

All custom mathematical and plotting operations were performed with Python 3.8.16. Scripts and additional information can be found on the GitHub repository of this project: https://github.com/Kanomble/eps_project. Software versions and all utilized packages and third-party tools are implemented in a public Docker image (kanomble/eps_project:1.2).

## Acknowledgements

We thank Katharina Grosche for excellent assistance with the carbohydrate analyses.

## Funding

This study was supported by the DFG (Project FR 3041/3-1). MK is supported by the DFG (Project 517563163). MP acknowledges support from the Cluster of Excellence on Plant Sciences (CEPLAS) funded by the Deutsche Forschungsgemeinschaft (DFG, German Research Foundation) under Germany’s Excellence Strategy–EXC 2048/1–Project ID: 390686111.

## Supplementary Tables and Figures

**Table S1.** Detailed information on the three free-living and three host associated *Curvibacter* strains used in the comparative genomics approach.

| Scientific Name | Taxonomic ID | Genome ID | Isolation Source<sup>1</sup> | Literature |
| --- | --- | --- | --- | --- |
| *Curvibacter lanceolatus*<br>ATCC 14669 | 1265504 | GCF_000381265.1 | Distilled water in<br>Vancouver, Canada | Ding & Yokota,<br>2004; Lyu et al.,<br>2024; Zhang et<br>al., 2017<sup>3</sup> |
| *Curvibacter gracilis* ATCC<br>BAA-807 | 1336244 | GCF_000518645.1 | Well-water in Osaka,<br>Japan | Ding & Yokota,<br>2004<sup>2</sup> |
| *Curvibacter delicatus* | 80879 | GCA_041639495.1 | Sediment of a pyritic acid<br>mine drainage in<br>Queensland, Australia | Ding & Yokota,<br>2004<sup>2</sup> |
| *Curvibacter*<br>Hmag1.1 | 1888168 | - | *Hydra magnipapillata* | - |
| *Curvibacter*<br>Hvul1 | 1888168 | - | *Hydra vulgaris* | - |
| *Curvibacter* sp.<br>AEP1.3 | 1844971 | GCF_002163715.1 | *Hydra vulgaris* AEP | - |

<sup>1</sup>The natural environment of the free-living strains differs. *C. lanceolatus* was isolated from distilled water in Vancouver, British Columbia, Canada. *C. gracilis* was isolated from well-water in Osaka, Japan (Ding & Yokota, 2004). *C. delicatus* was isolated from sediment, specifically from a pyritic acid mine drainage in Queensland, Australia.
<sup>2</sup>Beyond the initial description and characterization, there is no further information available on *C. delicatus* and *C. gracilis*.
<sup>3</sup>In contrast, *C. lanceolatus*, particularly the strain *C. lanceolatus* HJ-1, is recognized for its ability to induce carbonate mineralization.

**Table S2.** Comparison of OGs among *Curvibacter* AEP1.3 and its free-living and host-associated relatives.

| Other *Curvibacter* | *Curvibacter* sp. AEP1.3 | Unique OGs *Curvibacter* sp. AEP1.3 | Unique OGs other *Curvibacter* strain | $\Sigma$ OGs other *Curvibacter* strain |
| --- | --- | --- | --- | --- |
| *Curvibacter lanceolatus* | 2450 | 835 | 2307 | 4757 |
| *Curvibacter gracilis* | 2427 | 858 | 2300 | 4727 |
| *Curvibacter delicatus* | 1518 | 1767 | 328 | 1846 |
| *Curvibacter* Hmag1.1 | 3150 | 135 | 558 | 3708 |
| *Curvibacter* Hvul1 | 3152 | 133 | 587 | 3739 |

OGs = Orthogroups, $\Sigma$ OGs *Curvibacter* sp. AEP1.3 = 3285.
*C. lanceolatus* and *C. gracilis* have the highest (4757 and 4727) proportion of OGs, while *C. delicatus* has the fewest (1846) OGs. The three symbiotic strains possess a medium number of OGs (3285 - 3739). The number of OGs unique to *Curvibacter* AEP1.3 was lowest (133 and 135) in comparison with the other two *Hydra*-associated symbionts, indicating a close relationship between these strains. In contrast, in comparison with *C. delicatus*, *Curvibacter* AEP1.3 exhibited the highest number of unique OGs (1767), likely due to the smaller total OG count (1846) of *C. delicatus*. Conversely, the number of shared OGs was highest in comparison with both symbiotic strains (3150 and 3152), and lowest in comparison to *C. delicatus* (1518). The number of shared OGs between *Curvibacter* AEP1.3 and the two other free-living strains (2427 and 2450) was nearly half of their total OG count.

**Table S8.** Additional information on free-living *Curvibacter* strains.

| Scientific Name | Taxonomic ID | Genome ID | Isolation Source | Literature |
| --- | --- | --- | --- | --- |
| *Curvibacter* sp.<br>BRE_MAG_193 | 1888168 | GCA_038112835.1 | Estuarine biome, Brisbane, Australia | Fan et al., 2024 |
| *Curvibacter* sp.<br>BRE_MAG_257 | 1888168 | GCA_038111605.1 | Estuarine biome, Brisbane, Australia | Fan et al., 2024 |
| *Curvibacter* sp.<br>APW13 | 3077236 | GCF_032337135.1 | Artificial pond water, Dongguk University, South Korea | - |
| *Curvibacter* sp.<br>HBC28 | 3026421 | GCF_028649455.1 | Cyanobacterial bloom, South Korea | - |
| *Curvibacter* sp.<br>RS43 | 3026419 | GCF_028649435.1 | Cyanobacterial bloom, South Korea | - |
| *Curvibacter* sp.<br>AG1 | 1888168 | GCA_020621965.1 | Enrichment culture sampled from a low-nitrate anoxic artesian well, Altingen, Germany | Jakus et al., 2021 |
| *Curvibacter lanceolatus*<br>LaCa_Mag_1 | 86182 | GCF_019163435.1 | Lake Cadagno chemocline, Switzerland | Philippi et al., 2021 |
| *Curvibacter* sp.<br>PH2015_06S_60_13 | 1888168 | GCA_004293565.1 | River cyanobacterial mat, Eel River, California, USA | Bouma-Gregson et al., 2019, 2022 |
| *Curvibacter* sp.<br>AWTP1-17 | 1888168 | GCA_003987795.1 | Simulated distribution system annular reactor, Texas, USA | Kantor et al., 2019 |
| *Curvibacter* sp.<br>UBA10131 | 1888168 | GCA_003537855.1 | Groundwater metagenome, Queensland, Australia | Parks et al., 2018 |
| *Curvibacter* sp.<br>UBA7973 | 1888168 | GCA_003500955.1 | Marine metagenome, Queensland, Australia | Parks et al., 2018 |
| *Curvibacter* sp.<br>RIFCSPHIGHO2_12_FULL_63_18 | 1797748 | GCA_001795465.1 | Rifle well FP-101 under high O2 conditions, USA | Anantharaman, Brown, Burstein, et al., 2016; Anantharaman, Brown, Hug, et al., 2016; Kauffman et al., 2018; Zaremba-Niedzwiedzka et al., 2017 |
| *Curvibacter* sp.<br>GWA_63_95 | 1797746 | GCA_001795455.1 | Rifle well FP-101 under high O2 conditions, USA | Anantharaman, Brown, Burstein, et al., 2016; Anantharaman, Brown, Hug, et al., 2016; Kauffman et al., 2018; Zaremba-Niedzwiedzka et al., 2017 |
| *Curvibacter* sp.<br>PAE-UM | 1714344 | GCF_001432305.1 | River water, Nankai University, China | Ma et al., 2016 |

**Figure S1:**
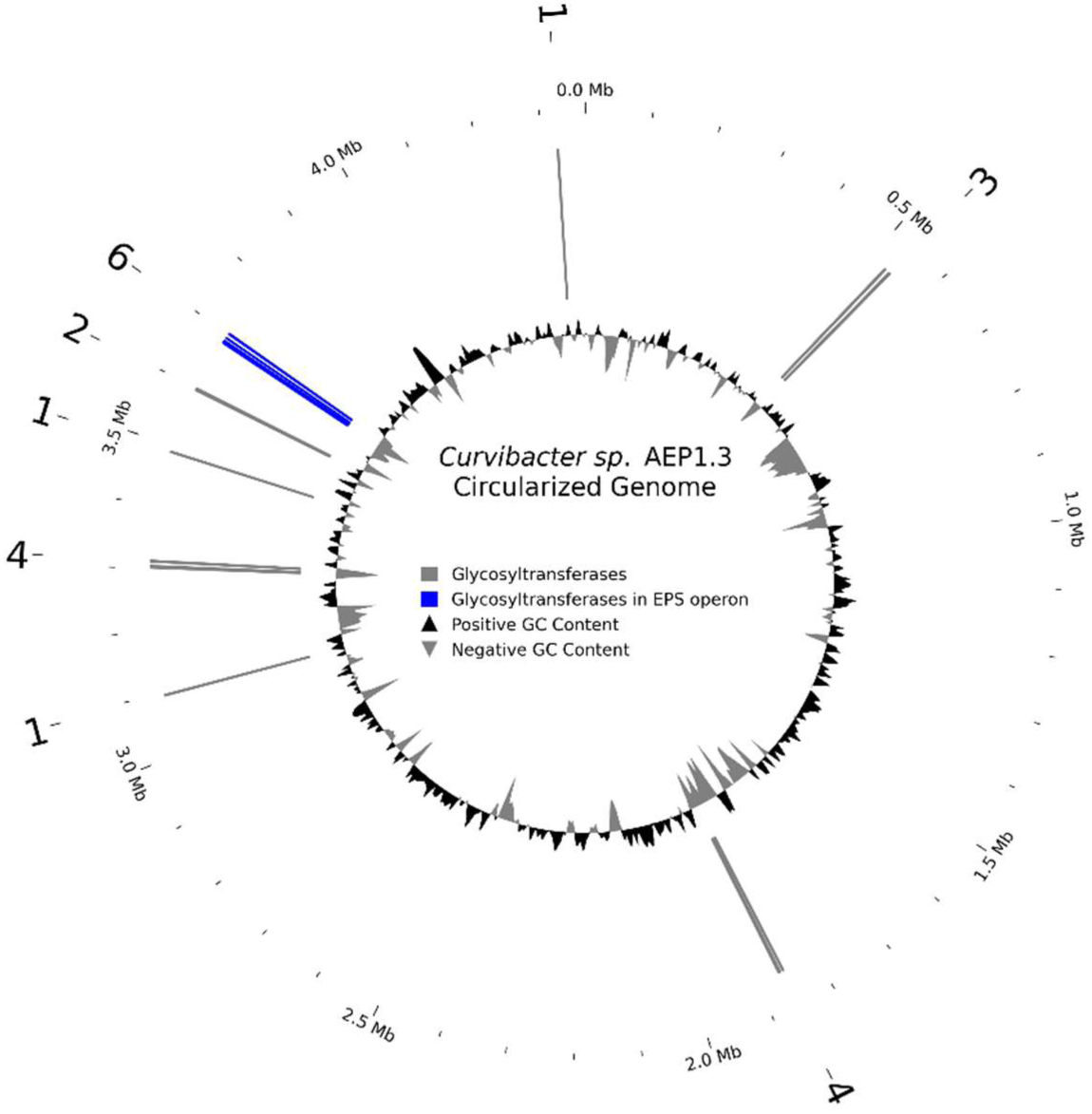
Distribution of glycosyltransferases genes within the genome of *Curvibacter* sp. AEP1.3. Potential glycosyltransferase sequences were obtained by screening coding DNA sequence annotations within the GenBank file of *Curvibacter* sp. AEP1.3. The innermost circle is composed of a density plot that shows the GC content of the respective genome regions. Positive GC-content refers to genomic regions where the GC content is higher than the genome-wide average, whereas negative GC content refers to regions where it is lower. The following circle, composed of gray and blue lines, indicates the locations of glycosyltransferases, with the associated numbers displayed in the outermost circle. Blue lines highlight the location of glycosyltransferases within (WcaJ and WecB) and near the EPS operon.

**Figure S2:**
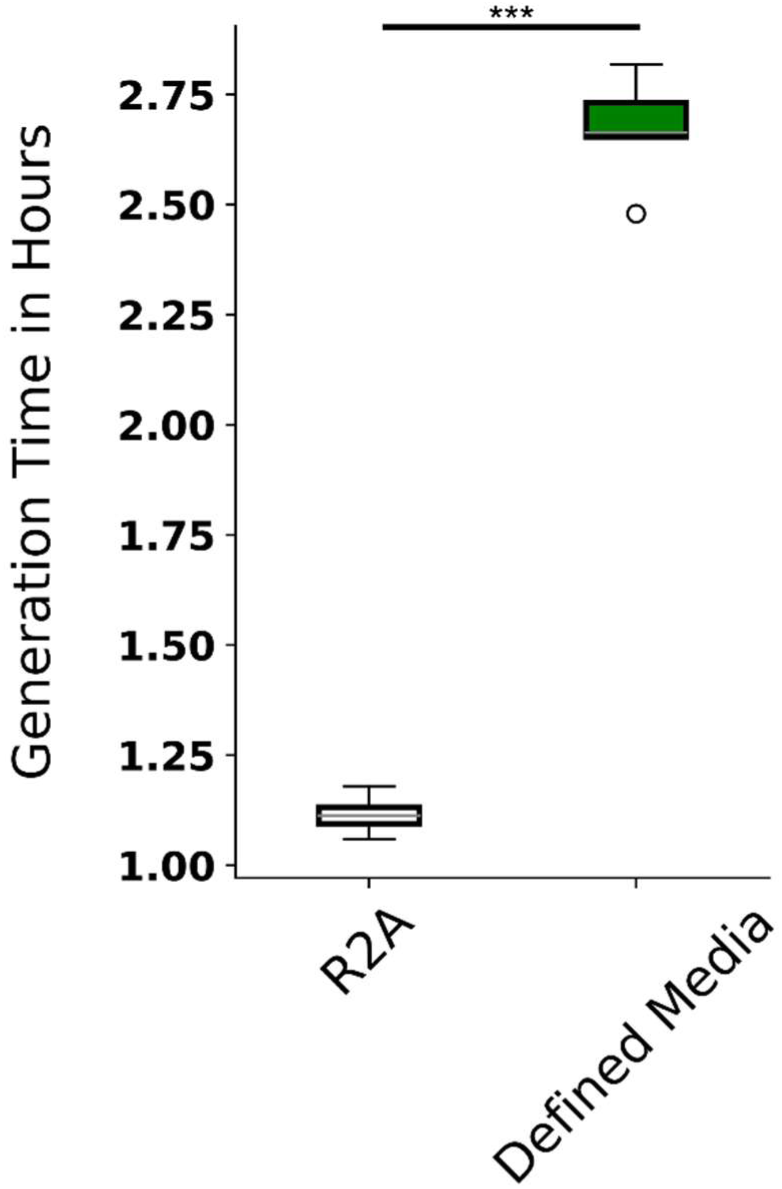
Generation time in hours of *Curvibacter* sp. AEP1.3 cultures in a CLARIOstar plate-reader at room temperature (N=6) inoculated in complex media (R2A) or defined media (Table 1). Statistical analyses were conducted using a one-way ANOVA, the comparison was conducted with Tukey’s HSD test to identify significant differences. Statistical significance is indicated by asterisks, with the following meanings: *p < 0.05, **p < 0.01, ***p < 0.001.

**Figure S3:**
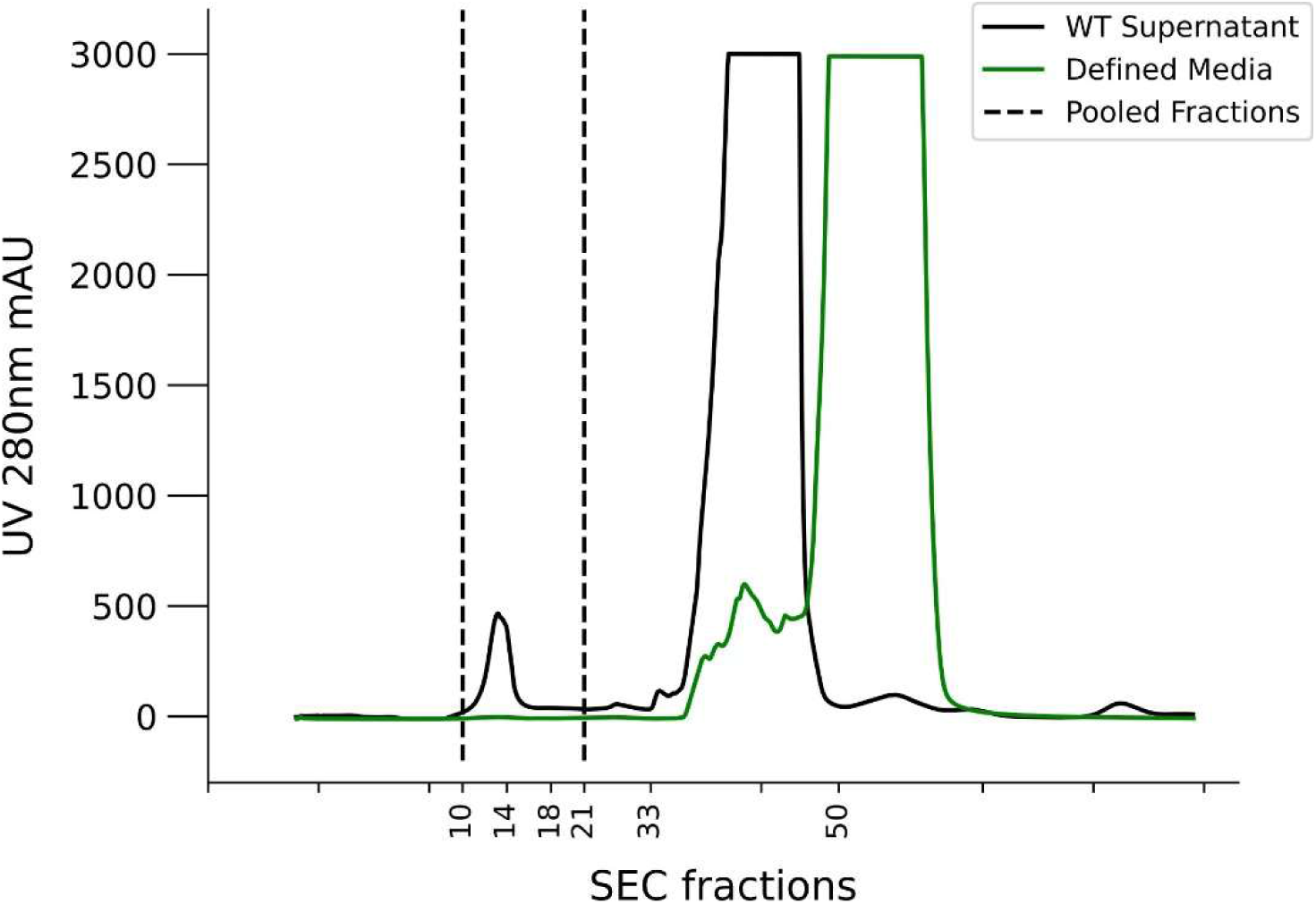
Chromatogram of the UV280 detector. Lyophilized, redissolved, cell-free supernatant of the *Curvibacter* sp. AEP1.3 wt supernatant (black) and the defined media as a negative control.

**Figure S4:**
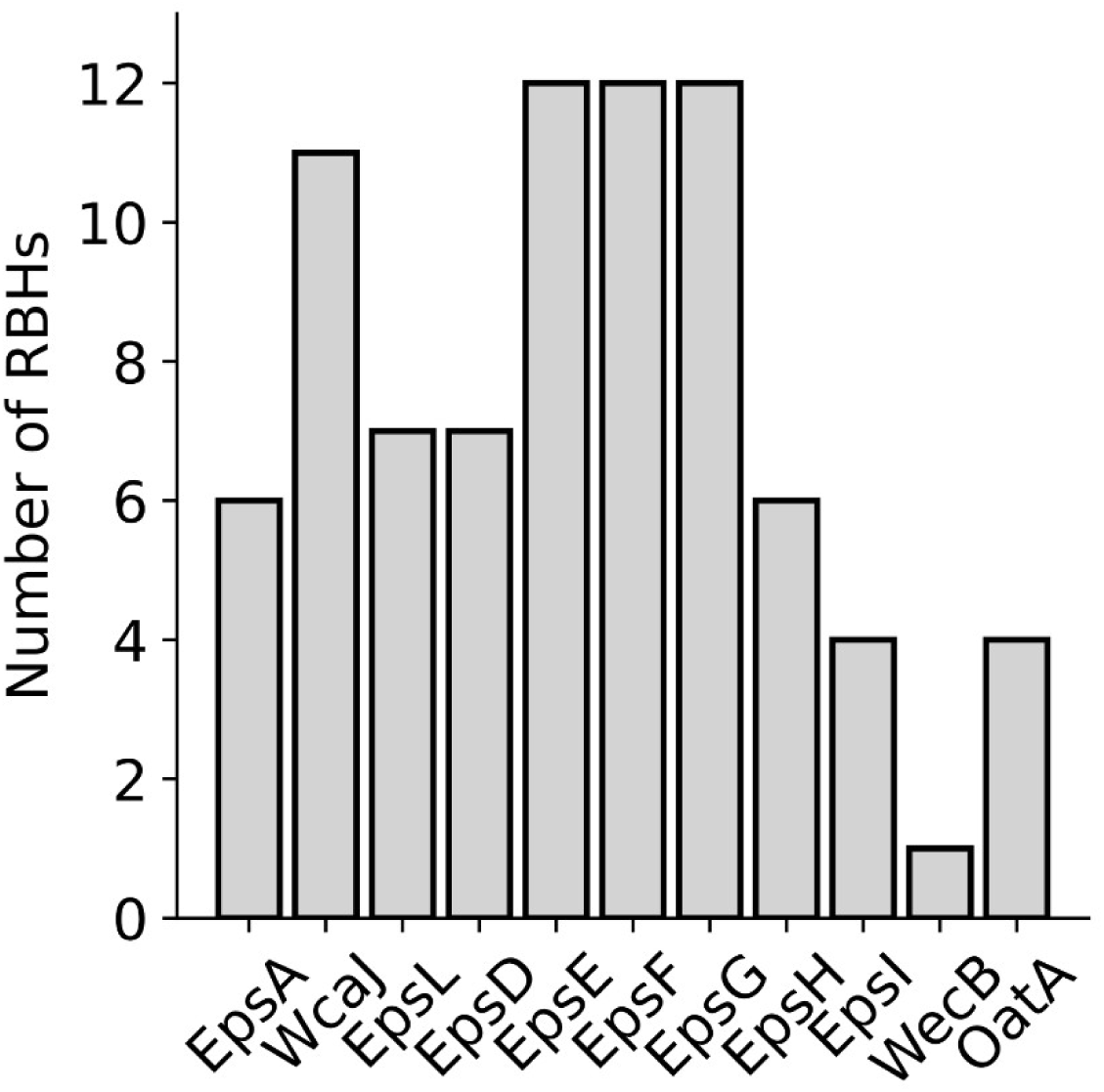
CATHI results of *Curvibacter* RBHs of the EPS operon. Most Reciprocal Best Hits (RBHs) can be observed for the EpsE, EpsF and EpsG protein sequences, therefore the synteny analysis (Figure 3D) has been applied based on the genomic location of those sequences. The complete CATHI results are summarized in supplementary table S9.

**Figure S5:**
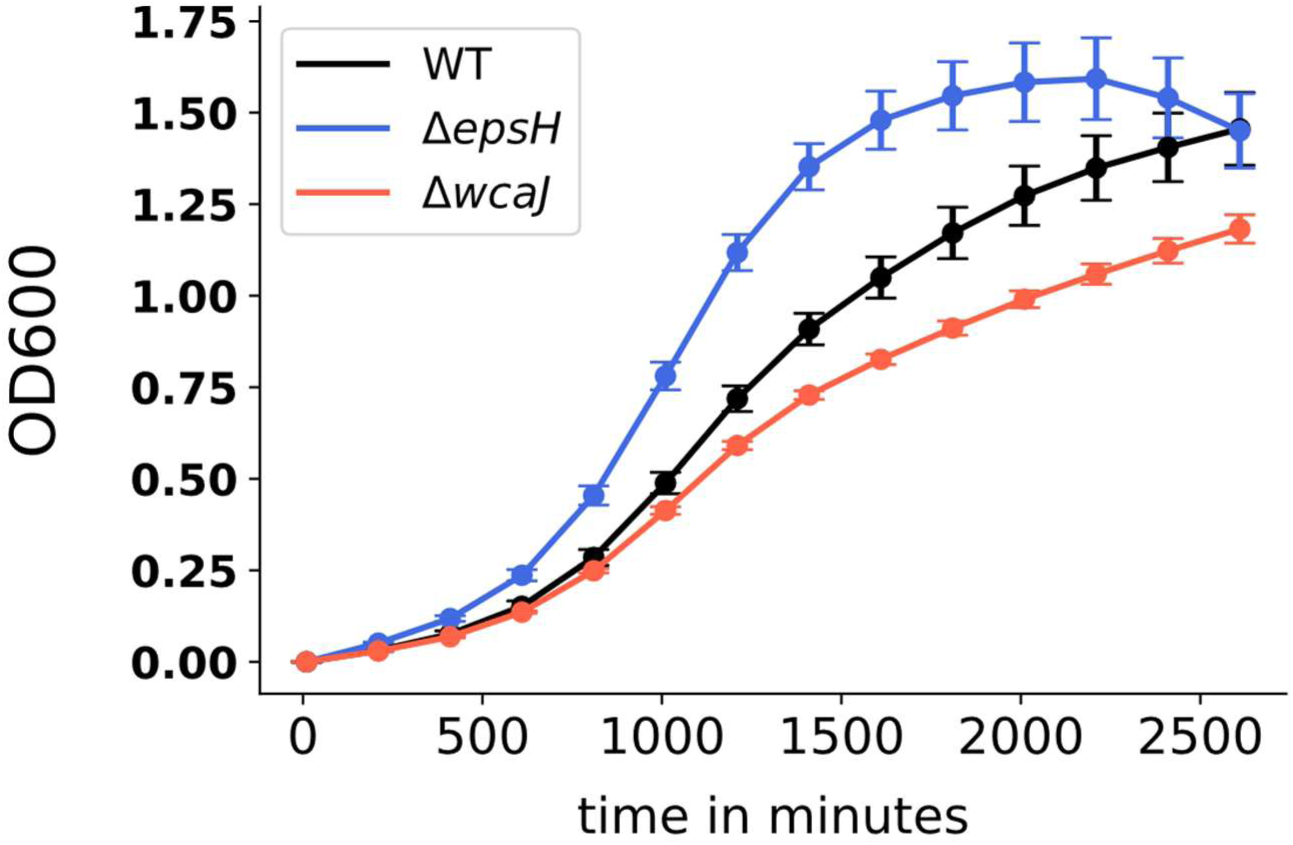
OD_600_ growth curves of *Curvibacter* sp. AEP1.3 wt, and the two mutant strains *ΔwcaJ* and *ΔepsH* (N=6). Growth was measured in a CLARIOstar machine at room temperature in defined media (Table 1). The growth curves correspond to the generation times displayed in Figure 5C.

**Figure S6:**
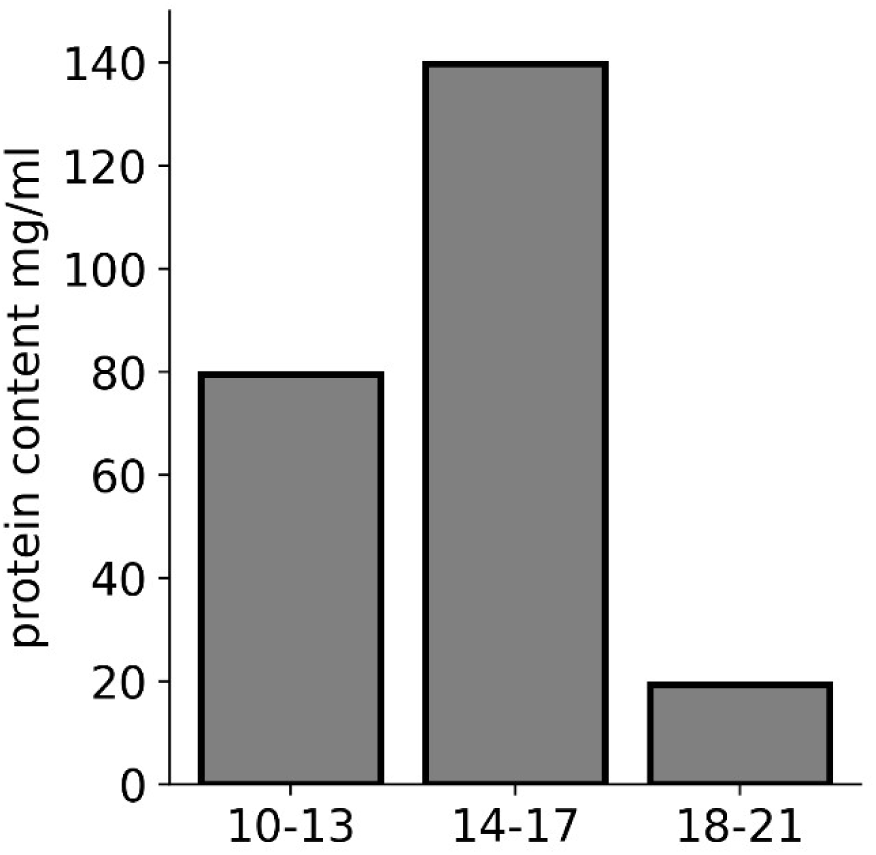
Bradford assay of the pooled SEC-fractions. Protein concentration within the high-molecular weight fractions of the SEC obtained from the lyophilized, redissolved, cell-free supernatant of the *Curvibacter* sp. AEP1.3 wt supernatant (N=3).

## Notes

https://github.com/Kanomble/eps_project

### Competing Interest Statement

The authors have declared no competing interest.

